# Dynamic changes in the urine proteome in two ovarian cancer rat models

**DOI:** 10.1101/604850

**Authors:** Yuqiu Li, Linpei Zhang, Wenshu Meng, Youhe Gao

## Abstract

Ovarian cancer is the most lethal gynecological malignancy in women, and it is likely to metastasize and has a poor prognosis. The early and reliable diagnosis and monitoring of ovarian cancer is very important. Without a homeostasis mechanism, urine can reflect early systemic changes in the body and has a great potential to be used for the early detection of cancer. This study tested whether early changes could be detected in two ovarian cancer rat models. Two rat models were established by either intraperitoneal (i.p.) or orthotopic (o.t.) injection of NuTu-19 ovarian cancer cells in female Fischer344 rats. Urine samples from ovarian cancer rats were collected at five time points during cancer development, and urinary proteins from the rats were profiled by liquid chromatography coupled with tandem mass spectrometry (LC-MS/MS). Compared with pre-injection samples, 49 differential proteins that have human orthologues were significantly changed in the orthotopically injected model. Among them, 24 of the differential proteins have previously been reported to be associated with ovarian cancer, six of which were reported to be biomarkers of ovarian cancer. On the 7th day after orthotopic injection, four differential proteins (APOA1, OX2G, CHMP5, HEXB) were identified before obvious metastases appeared. In the intraperitoneal injection model, 76 differential proteins were changed during the course of ovarian cancer development. The results show that urine proteins could enable the early detection and monitoring of ovarian cancer progression and could lay a foundation for further exploration of the biomarkers of ovarian cancer.

## 1. Introduction

Ovarian cancer (OC) is the most lethal gynecological malignancy and it is the fifth leading cause of cancer death in women^[1]^, mainly because OC can spread to intraperitoneal tissues, such as the omentum in the peritoneal cavity, by the time of diagnosis^[2]^. The strongest risk factors for this disease are an advanced age and a family history of ovarian cancer. Epithelial ovarian cancer (EOC) is the most common ovarian malignancy and comprises 90% of all ovarian cancers^[3]^. More than 75% of affected women ovarian carcinomas are not diagnosed until a late stage (stage III or IV) because early-stage disease is not obvious and early symptoms, as well as symptoms of late-stage disease, are nonspecific. Pelvic or abdominal pain, a feeling of pelvic mass and abdominal swelling are the main clinical symptoms, but these are not specific to OC^[4]^. Currently, women who have symptoms consistent with ovarian cancer usually undergo a physical examination, transvaginal ultrasonography, and measurement of biomarkers such as CA125, but imaging and serum biomarkers tests are of limited value in the early diagnosis of the ovarian carcinomas^[5]^. The treatment of ovarian cancer usually involves surgery and intraperitoneal and intravenous chemotherapy^[6]^. At present, patients are typically diagnosed at the advanced stage when the cancer has disseminated within the peritoneal cavity, and complete surgical removal is impossible^[3]^. An early diagnosis is important, since the five-year survival rate for women diagnosed with stage I epithelial ovarian cancer is over 90% compared with a 25% five-year survival rate for stages III and IV^[4]^. An early diagnosis when tumors are small and still confined to the ovaries is the most important prognostic factor^[6]^. Therefore, early detection and cancer screening have the potential to decrease the mortality and morbidity of this cancer. If diagnosed earlier, effective measures would be used to prevent the development of ovarian cancer. Because of the variety of the pathology and an unclear mechanism and etiology, there are currently no tumor markers with both high specificity and sensitivity that could be applied to the early clinical diagnosis of ovarian cancer^[2]^. Thus, the problem of how to detect and diagnose ovarian cancer with better sensitivity in the early stages need to be urgently addressed. A novel systematic method with high sensitivity and specificity for the early diagnosis of OC and new ovarian cancer markers need to be identified.

Biomarkers are measurable changes related to physiological or pathological processes^[7]^. Without a homeostasis mechanism, urine can sensitively reflect early changes in the body and has the potential to be used for the early detection of cancer. Technology advances in urinary proteomics have allowed it to play a significant role in tumor biomarker discovery^[8]^. Candidate biomarkers of different diseases have been detected in urine, and some of these biomarkers even perform better in urine than in serum [9]. We investigated whether the urinary proteome could reflect early changes in and the development of ovarian cancer. The urine accommodates a variety of changes, and urinary proteins were found to be associated with gender, age, diet, exercise, hormone status and other physiological conditions^[10]^. Previous studies have identified several biomarkers in the clinical urine samples of women with ovarian cancer, including HE4^[11]^, eosinophil-derived neurotoxin, a fragment of osteopontin^[12]^, mesothelin^[9]^ and Bcl-2^[13]^. Most of the collected clinical samples were from advanced cancer stages. Using clinical samples to find early biomarkers requires a large number of early-stage samples to balance the population and individual differences, which requires much time and carries a high cost. However, using simple animal models to study urine biomarkers can reduce the sample complexity, and it is also possible to monitor the process of disease development and observe related physiological changes in the early stages of the disease for early diagnosis^[14]^. Experimental models are crucial to understanding the biological factors that influence the phenotypic characteristics of the disease^[15]^. A research model that enables a focus on early-stage disease would prove beneficial. Biomarker research using animal models has been conducted to study various diseases, including obstructive nephropathy^[16]^, pulmonary fibrosis^[17]^, hepatic fibrosis^[18]^, myocarditis^[19]^ and glomerular diseases^[20]^, chronic pancreatitis^[21]^, Alzheimer’s disease^[22]^, and subcutaneous tumors ^[23]^. These studies have found potential urinary biomarkers in the early stages of diseases by applying proteomic technology.

Epithelial ovarian cancer has four common subtypes: serous, endometrioid, clear cell, and mucinous carcinoma^[24]^. NuTu-19 is a surface epithelial cell line derived from serous epithelial ovarian cancer of Fischer 344 (F344) rats that underwent spontaneous malignant transformation in vitro. It was reported that the NuTu-19 cell line was injected into naive, immunocompetent F344 rats to determine tumor growth and animal survival^[25]^. This was a spontaneously arising, nonimmunogenic, experimental animal model of epithelial ovarian cancer that has the ability to mimic the development of ovarian cancer.

Ovarian cancer is more likely to metastasize via intraperitoneal dissemination than by hematogenous metastasis. Intraperitoneal spread appears to be an early event, which is why these cancers are only rarely detected at an early stage^[26]^. The physiological movement of ovarian cancer cells from the primary tumor to the peritoneum and omentum, as well as the direct expansion of tumor lesions to adjacent organs, has widely been recognized as the most common metastatic route of epithelial ovarian cancer^[27]^. One model utilizes the intraperitoneal injection of NuTu-19 cells in immunocompetent F344 rats. This is an inexpensive, reproducible and efficient preclinical model in which to study ovarian peritoneal carcinomatosis^[28]^. NuTu-19 cells injected intraperitoneally can grow progressively as numerous serosal nodules, exhibit local tissue invasion and form malignant ascites in a manner typical of human ovarian epithelial carcinomas^[25]^. This syngeneic model, which directly introduces the ovarian cancer cells into the peritoneum immediately produces extrapelvic extension that quickly creates metastases that resemble the spreading found in the late stages of the disease^[29]^. In addition, a previous study showed used a modified approach to combine surgical orthotopic implantation techniques with the syngeneic NuTu-19 cells. This created a novel model that initiates a primary lesion within the inherent microenvironment to mimic the early stages and natural progression of primary ovarian cancer^[30]^. The orthotopic model is more similar to the clinical process of human ovarian cancer. It offers a clinically relevant alternative for primary ovarian cancer research that enables the investigation of an early diagnosis for this disease process.

In the current study, we attempted to answer the following questions: 1) Can urine proteins sensitively reflect the disease progression at various stages of ovarian peritoneal carcinomatosis or abdominal metastasis of cancer? 2) Can urinary proteins sensitively reflect early changes in early-stage primary ovarian cancer, and how could the urinary biomarkers before metastasis of the ovarian cancer be characterized for early diagnosis? We analyzed the urinary proteome in two ovarian cancer rat models by using LC-MS/MS. The workflow of the urinary proteomics analysis is shown in Fig. 1. Our aim was to identify differential urinary proteins associated with disease progression and early lesions and to provide some clues for monitoring the disease and early diagnosis of ovarian cancer.

**Fig. 1.**
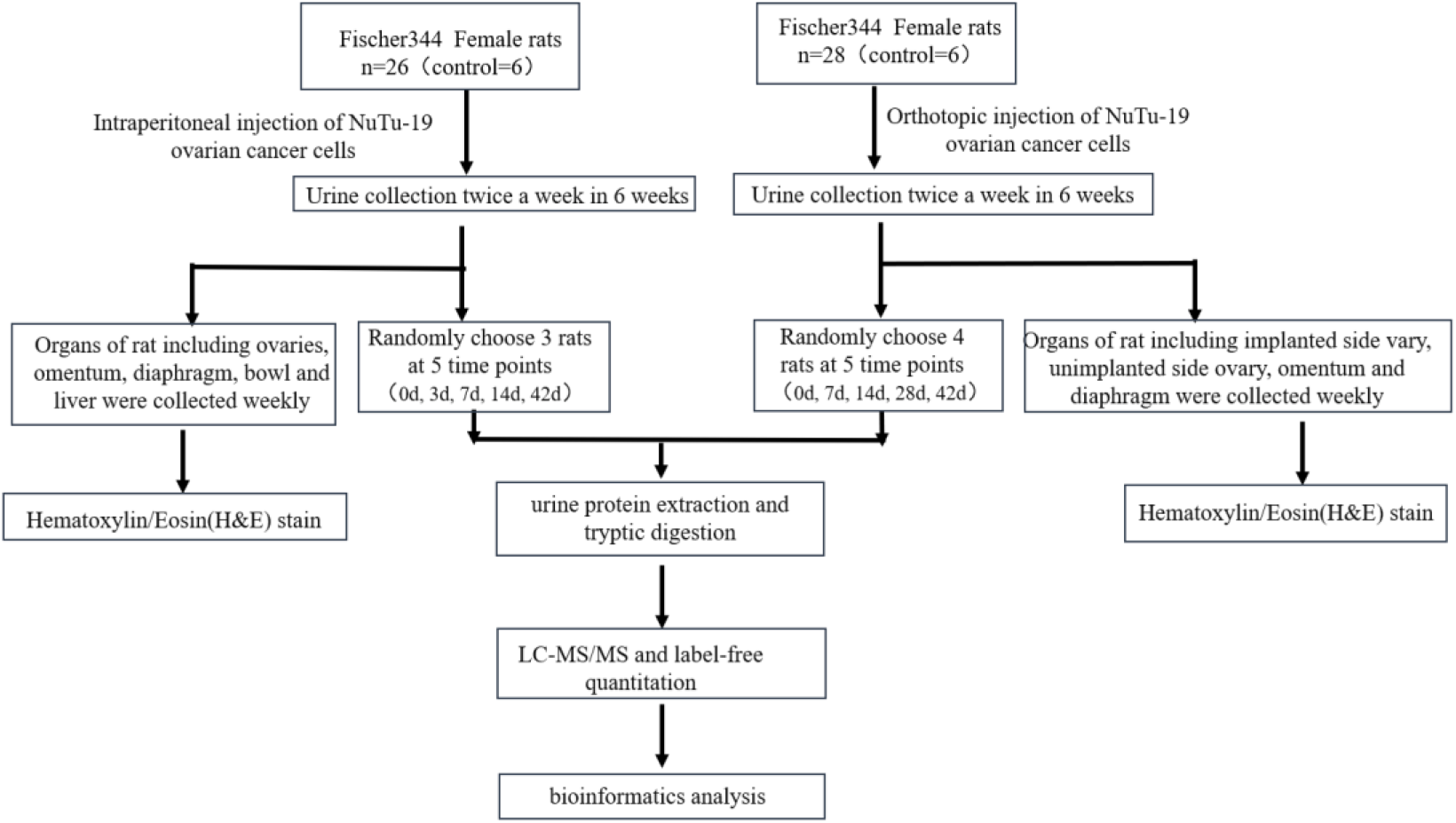
The urinary proteome analysis workflow in ovarian cancer rats. Urine samples were collected in each phase, and histological analyses of the ovaries and other organs were conducted. Urinary proteins were identified at five time points by LC-MS/MS.

## 2. Materials and methods

### 2.1 NuTu-19 cell line and establishment of ovarian cancer rat models

The cell line NuTu-19 was derived from epithelial ovarian cancer of F344 female rats (type: MXC290, brand: ATCC) and was purchased from Shanghai Meixuan Biotechnology Co., Ltd. (Meixuan, China). The cells were maintained in Roswell Park Memorial Institute (RPMI-1640) media (Corning, USA) supplemented with 1% penicillin–streptomycin and 10% fetal bovine serum (Gibco, USA) in an atmosphere of 95% air and 5% CO2 at 37 °C and passaged when they reached approximately 80% confluency. The cells were subcultured with trypsin ethylene-diaminetetraacetic acid (EDTA) every other day at split ratios of 1:3. The NuTu-19 cells were harvested using trypsin–EDTA to generate a single-cell suspension using physiological saline; these cells were stained with trypan blue (BioWhittaker) and counted using a hemocytometer (American Optical, Buffalo, NY) to test cell viability and determine the viable concentration.

Specific pathogen-free 6–8-week-old (100-120 g) female Fischer 344 (F344) rats (purchased from Vital River Laboratory Animal Technology Co., Ltd., China) were housed at a constant temperature and relative humidity, with a standard 12 h light/12 h dark cycle and in a standard environment (room temperature 22 ± 1 °C, humidity 65%-70%). The animals were acclimated to the animal room for 1 week before experiments. The animal license number was SCXK (Beijing) 2016-0011. The rats were given ad libitum access to feed and water. The experiment in this study was approved by the Institutional Animal Care Use & Welfare Committee of the Institute of Basic Medical Sciences, Peking Union Medical College (Animal Welfare Assurance Number: ACUC-A02-2014-007).

The NuTu-19 ovarian cancer cell line with intraperitoneal injection to induce an ovarian peritoneal carcinomatosis rat model was established as follows^[25]^: A total of 26 female F344 rats from the first batch were randomly divided into two groups: the control group (n=6) and the experimental group (n=20). Rats in the experimental group were intraperitoneally injected with 2.5×10^6^ NuTu-19 cells in a total volume of 0.5 ml by 1 ml sterile syringe (BD, China). At the same time, rats in the control group were given an intraperitoneal injection of 0.5 ml normal saline. The injection site was in the right lower quadrant. The rats were observed daily and weighed weekly.

The orthotopic injection of NuTu-19 cells induced the primary ovarian cancer rat model that was established by the epithelial ovarian cancer line as follows^[30]^: A total of 28 female F344 rats from the second batch were randomly divided into two groups: the control group (n=6) and the experimental group (n=22). All rats were anesthetized by intraperitoneal injection of 20 mg/kg body weight of 2% pentobarbital sodium (Sinopharm Chemical Reagent Co., Ltd. China). The anesthetized F344 rats were fixed on the rat plate. After local disinfection, all subsequent procedures were conducted under aseptic conditions. In the middle of the lower abdomen and 1 cm above the pubic symphysis, the scalpel was longitudinally incised approximately 1.5 cm. When the right side ovary was fully exposed, carefully stabilized with surgical instruments, and 5 μl of suspension was injected with a microinjection needle (Shanghai Gaoge Company, China) under the right ovarian capsule. In the experimental group, using an orthotopic injection technique, the 2×10^4^ NuTu-19 cells in a total volume of 5 μl were inoculated just below the epithelial bursa surrounding of the ovary. In the control group, the rats were given 5 μl of normal saline solution by using an orthotopic injection technique. After the animals were awake, they were returned to the cage and each rat was intraperitoneally injected with 0.25 ml of 80,000 units of penicillin and kept in a clean environment. These rats were observed and weighed weekly.

### 2.2 Histological analysis of ovarian cancer rats

In the intraperitoneal (i.p.) model, nine rats in the experimental group and three rats in the control group were randomly sacrificed per week after injection. The organs (ovaries, omentum, diaphragm, bowl and liver) of the experimental rats and control rats were harvested on the 7th, 14th, 28th and 42nd day after the intraperitoneal injection. On the 42nd day, all rats were sacrificed and the abdominal organs were taken for pathology. In the orthotopic (o.t.) model, some rats in the experimental group and rats in the control group were randomly sacrificed weekly after injection. On the 7th, 14th, 28th and 42nd day after the orthotopic injection of cancer cells, various organs including implanted side ovary, unimplanted side ovary, omentum and diaphragm of rats in the control group and the experimental group were collected. All rats were sacrificed on the 42nd day, and the organs were harvested for histological and morphometric analysis. For histopathology, the abdominal organs were fixed in formalin (10%) and embedded in paraffin. The histopathological lesions and invasion were evaluated with H&E staining.

### 2.3 Urine collection and sample preparation

The urine of the experimental group was collected as a self-control urine sample one day before the model was established. Urine samples were collected from the experimental group twice per week for six weeks. Without any special treatment, the rats were individually placed in metabolic cages overnight for 12 h to collect urine. During the collection, no water or food was supplied to avoid urine contamination. After collection, the urine samples were stored at −80 °C. Urine samples from the experimental and control groups were centrifuged at 12,000g for 30 min at 4 °C to remove impurities and large cell debris. After removing the pellets, three volumes of prechilled ethanol were added and incubated at −20 °C for 2 h. After centrifugation, lysis buffer (8 mol/L urea, 2 mol/L thiourea, 50 mmol/L Tris, and 25 mmol/L DTT) was used to dissolve the pellets, which were then centrifuged at 12.0 g for 30 min at 4 °C. The supernatant containing the resulting protein extract was placed into a new tube. The protein concentration was determined by the Bradford assay.

The urinary protein samples at different time points were digested using the filter-aided sample preparation (FASP) method ^[31]^. A 100 μg aliquot of protein was transferred into a 10 kDa filter device (Pall, Port Washington, NY, USA). After sequentially washing with UA (8 mol/L urea, 0.1 mol/L Tris-HCl, pH 8.5) and 25 mmol/L NH4HCO3. Briefly, the proteins were denatured by dithiothreitol (DTT, Sigma), alkylated by iodoacetamide (IAA, Sigma), and the samples were centrifuged at 14,000 g for 30 min at 18 °C, followed by washing once with UA and three times with 25 mmol/L NH4HCO3. The samples were digested with trypsin (Promega, USA) (enzyme-to-protein ratio of 1:50) at 37 °C overnight. The digested peptides were desalted using Oasis HLB cartridges (Waters, Milford, MA) and then dried by vacuum evaporation (Thermo Fisher Scientific, Bremen, Germany) and stored at −80 °C.

### 2.4 LC-MS/MS analysis

The peptide samples resulting from the above digestion were separated by an EASY-nLC 1200 HPLC system (Thermo Fisher Scientific, USA). First, the digested peptides were dissolved in 0.1% formic acid to a concentration of 0.5 μg/μl, and the BCA assay was used to determine the peptide concentration. Next, 1 μg of peptides from an individual sample was loaded onto a trap column (Acclaim PepMap®100, 75 μm×100 mm, 2 μm, Nano Viper C18). The elution gradient was 5%-40% mobile phase B (mobile phase A: 0.1% formic acid; mobile phase B: 89.9% acetonitrile, column flow rate 0.3 μl/min) over 60 min. The peptides were analyzed by an Orbitrap Fusion Lumos Tribrid mass spectrometer (Thermo Fisher Scientific, USA)^[32]^. MS data were acquired in data-dependent acquisition mode using the following parameters. Survey MS scans were collected by the Orbitrap in a 350–1550 m/z range and MS scans at a resolution of 120,000. For the MS/MS scan, the resolution was set at 30,000 screening in Orbitrap, and the HCD collision energy was 30, charge-state, dynamic exclusion (exclusion, duration 30 sec). Fifteen urine samples from three randomly selected i.p. injection-induced OC rats at five time points (on days 0, 3, 7, 14 and 42) were chosen for MS analysis. Additionally, twenty urine samples from three randomly selected o.t. injection-induced primary OC rats at five time points (on days 0, 7, 14, 28 and 42) were chosen for MS analysis. Two technical replicate analyses were performed for each urine sample.

### 2.5 Label-free proteome quantification and data processing

All MS data were processed using Proteome Discoverer (PD) software (version 2.4.1, Matrix Science, London, UK), and then MS data were searched using Mascot Daemon software (version 2.5.1, Matrix Science, UK) with the SwissProt_2017_02 database (taxonomy: Rattus; containing 7992 sequences). The search parameters were set as follows: The parent ion tolerance was set to 10 ppm, and the fragment ion mass tolerance was set to 0.05 Da; the carbamidomethyl of cysteine was set as a fixed modification; the oxidation of methionine was considered a variable modification; trypsin digestion was selected; and two sites of leaky cutting were allowed. The peptide and protein identifications were further validated using Scaffold (version 4.4.6, Proteome Software Inc., Portland, Oregon, USA). The setting conditions were as follows: Protein identification required a confidence score ≥ 95%; protein identifications were accepted at an FDR less than 1.0%; each protein contained at least 2 unique peptides; and spectral counting was used to compare protein abundance between different time points^[33,34]^.

In the i.p.-induced OC rat model, the proteins identified on the 3rd, 7th, 14th and 42nd day were compared with those from day 0. In the o.t.-induced OC rat model, the proteins identified on days 7, 14, 28 and 42 were compared with those from day 0. The changed urinary proteins were defined as those with a p value < 0.05 by a two-sided, unpaired t-test and a fold change >1.5 in the experimental groups when separately compared with self-control samples. Briefly, the differential proteins were selected with the following criteria: Each protein contained at least 2 unique peptides; fold change in the increased group ≥1.5 and fold change in the decreased group ≤0.67; and P < 0.05 by independent sample t-test. Moreover, the differential proteins at each time point showed a consistent trend in each experimental rat at that time point. Group differences resulting in P < 0.05 were identified as statistically significant. The statistical analysis was performed with GraphPad Prism version 5.0 (GraphPad, San Diego, CA).

### 2.6 Functional analysis of differential proteins

DAVID 6.8 (https://david.ncifcrf.gov/) was used to characterize the functional annotation of the differential urinary proteins identified at different time points in the i.p. and o.t. injection models. The detailed annotation included the biological process. All identified proteins were also analyzed by Ingenuity Pathway Analysis (IPA) software to enable a detailed annotation regarding canonical pathways.

## 3. Results and discussion

### 3.1.1 Characterization of the intraperitoneal injection ovarian cancer rat model

Rats in the control group had normal body weights. The body weight changes of the rats are shown in Fig. 2. After the 35th day after cell injection, the body weights of the intraperitoneal injection group were increased compared to those of the control group, and there was a significant difference. The weight change of the experimental rats may have been due to the presence of bloody ascites in the experimental group in the advanced stages of the disease. On the 35th day after intraperitoneal injection, some experimental rats of the i.p. injection group showed a reduced consumption of drinking water, rough hair, and the gradual appearance of abdominal distension. On the 42nd day after intraperitoneal injection, there were obvious symptoms, such as frog-like abdomen distension, pale auricle, and anemia. Upon dissection, numerous tumor nodules were found in the abdominal cavity, along with omental contraction. The surfaces of the ovary, diaphragm, liver and bowl were covered with cancer cells, and a large amount of malignant bloody ascites was present.

**Fig. 2.**
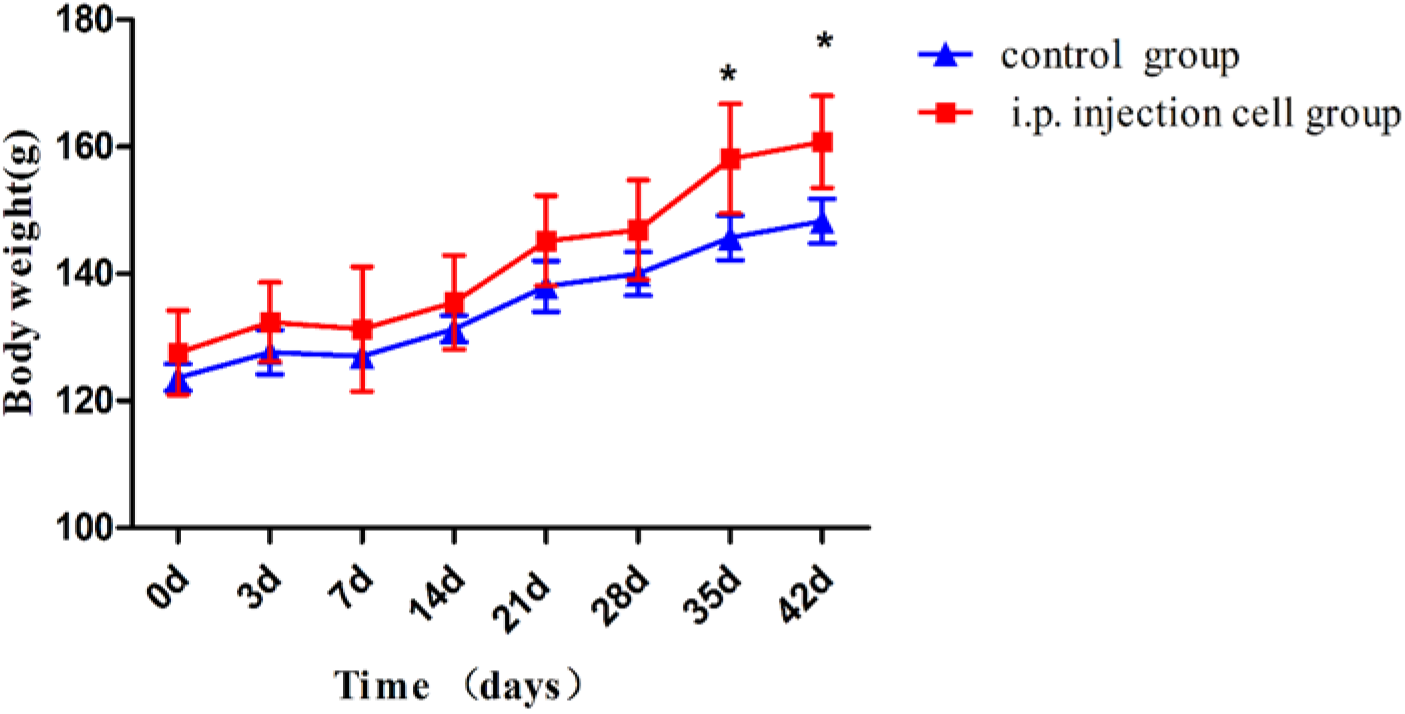
Changes in rat body weight during the intraperitoneally injected NuTu-19 cell experiment. The red line indicates the control group, and the blue line indicates the intraperitoneal NuTu-19 ovarian cancer cell group. * indicates P < 0.05.

Histopathological examination (H&E staining) of the ovaries and other abdominal organs of the rats were performed to determine the severity of the disease. The results of HE staining showed that there was no tumor invasion observed in the abdominal organs of the i.p. rats compared with the control group on the 7th day after the intraperitoneal injection (Fig. 3). At 14 days after i.p. injection, tumor infiltration first appeared in the omentum, which is a tissue attached to the greater curvature of the stomach and is the most common site for EOC metastatic spread ^[35]^. As the disease progressed, a greater extent of the omentum was invaded by cancer cells (Fig. 3A). Twenty-eight days after the injection of tumor cells, the ovary was invaded with cancer cells on the oviduct side (Fig. 3B), and the diaphragm was also invaded by cancer cells (Fig. 3C). On the 42nd day after injection, the bowel and liver tissue began to be infiltrated by the parenchyma (Fig. 3E-3F). At the same time, the peritoneal organs such as the omentum, diaphragm, and ovaries were seriously invaded by tumor cells. Microscopically, large numbers of tumor cells were observed in the tissues; the tumor cells were disordered, with an abnormal size and shape, and the cancer tissue appear poorly differentiated, which was consistent with the pathological features of ovarian cancer found in previous studies ^[30]^. These pathological changes revealed the successful induction of a peritoneal carcinomatosis rat model via the intraperitoneal injection of NuTu-19 cells.

**Fig. 3.**
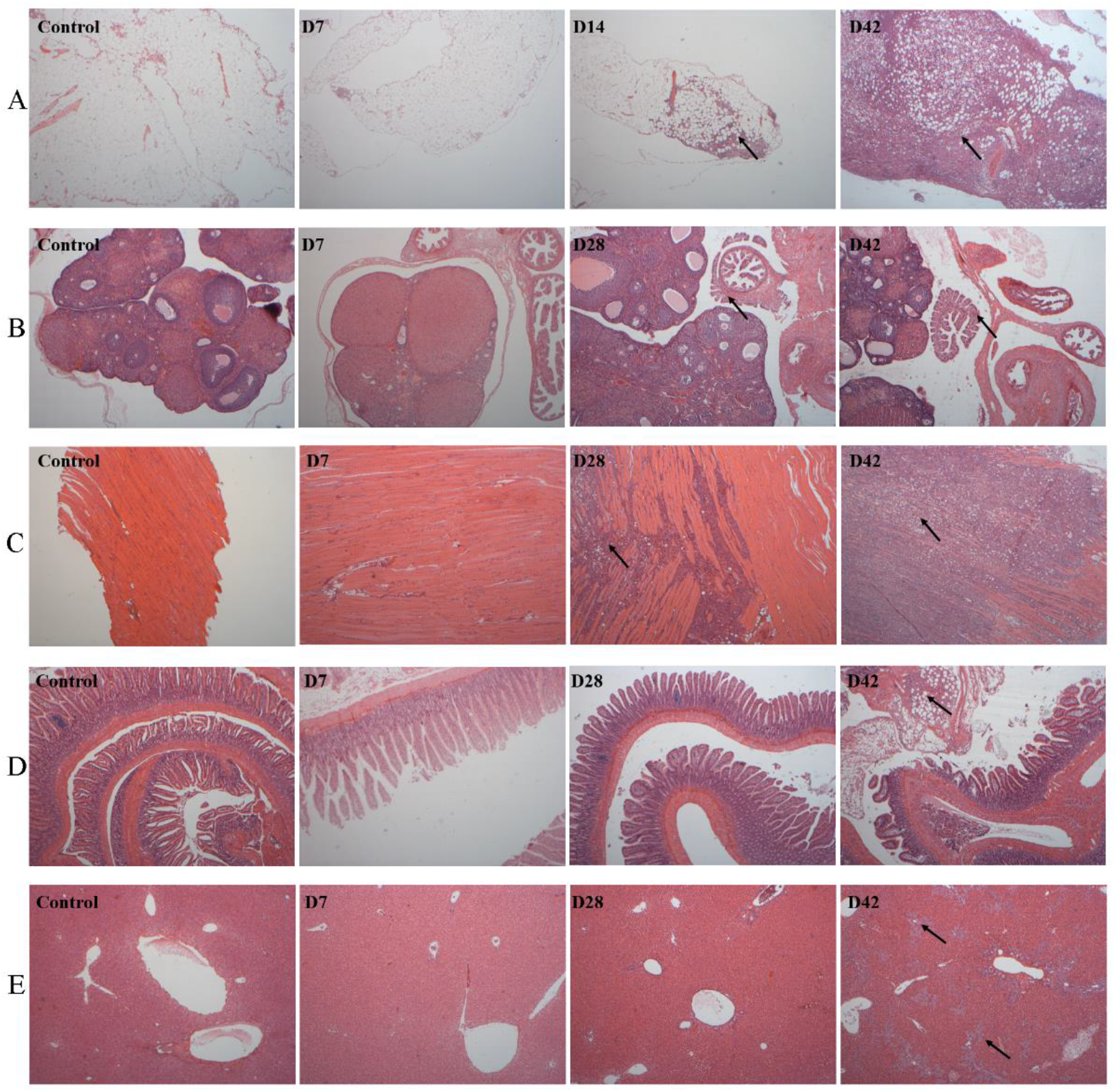
Histopathological characterization of organs in the NuTu-19 cell intraperitoneal injection rat model (40×). Organs were collected and prepared for microscopic examination using hematoxylin & eosin (H&E) stain. Metastases were identified in the A) omentum, B) ovaries, C) diaphragm, D) bowel, and F) liver. Each black arrow denotes the location of the metastasis in the tissue section.

### 3.1.2 Urine proteome changes in NuTu-19 cell intraperitoneal-injection OC model

To investigate how the urine proteome changes after the intraperitoneal injection of NuTu-19 ovarian cancer cells, urine samples collected at five time points (days 0, 3, 7, 14 and 42) from three experimental rats were analyzed by LC-MS/MS. In total, 577 urinary proteins were identified and are listed in Table S1. Among these, 76 differential proteins that had human orthologues were evaluated and were found to be significantly changed in the three rats (fold change ≥1.5 or ≤0.67, P < 0.05; Table 1). On the 3rd day, nineteen differential proteins were identified, two of which increased and seventeen of which decreased. On the 7th day, twenty-two differential proteins were identified, thirteen of which increased and nine of which decreased. On the 14th day, eleven differential proteins were identified, six of which increased and five of which decreased. On the 42nd day, thirty-seven differential proteins were identified, three of which increased and thirty-four of which decreased. Details of the differential proteins are presented in Table S2. Thirteen differential proteins were repeatedly identified at no less than one time point (Fig. 4), and the trend of the differential proteins was consistent at each time point in each rat. On the 3rd and 7th day, no obvious metastasis of the cancer tissue was found in the pathology of the abdominal organs, and 19 and 22 differential proteins were respectively identified in the urine. Among the 76 differential proteins, 53 are reported to be associated with cancer. Twenty-three proteins have been reported to be closely related to ovarian cancer and related diseases. Nine proteins have been reported as biomarkers of ovarian cancer. The urine protein profile also changed significantly as the NuTu-19 cell ovarian cancer cell proliferation in the abdominal cavity progressed.

**Fig. 4.**
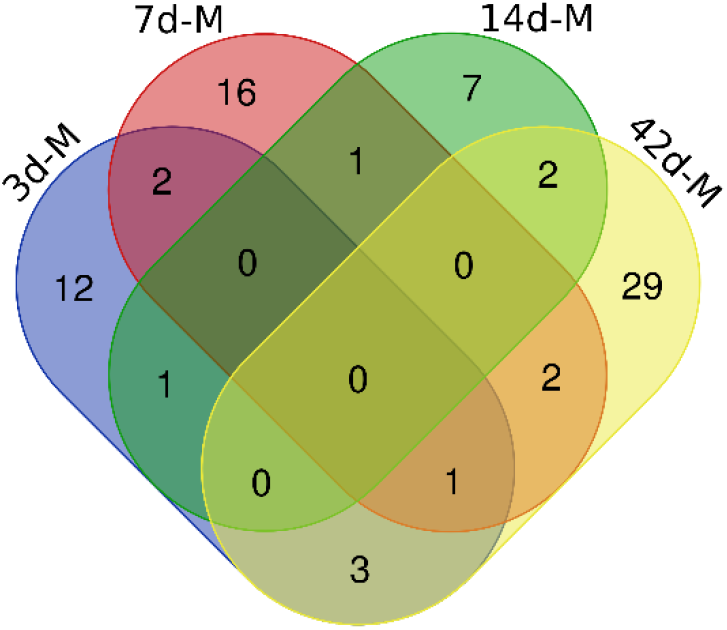
Venn diagram of the differential urinary proteins identified on days 3, 7, 14 and 42 after intraperitoneal injection of NuTu-19 cells.

**Table 1.** Differential urinary proteins in intraperitoneal injection NuTu-19 cell OC rats.

| Accession | Protein name | Trend | Fold change |  |  |  |  | P value | Reported to be related to ovarian cancer |
| --- | --- | --- | --- | --- | --- | --- | --- | --- | --- |
|  |  |  | 0d | 3d | 7d | 14d | 42d |  |  |
| Q1WIM1 | Cell adhesion molecule 4(CADM4) | ↑ | 1 | 1.77 | — | — | — | 3.94E-02 | tissue, serum<sup>[36]</sup> |
| P06757 | Alcohol dehydrogenase 1 (ADH1) | ↑ | 1 | 1.54 | 2.04 | — | — | 3.14E-02 | tissue<sup>[37]</sup> |
| Q6P734 | Plasma protease C1 inhibitor (IC1) | ↓ | 1 | 0.66 | — | 0.62 | — | 2.11E-03 |  |
| O70535 | Leukemia inhibitory factor receptor (LIFR) | ↓ | 1 | 0.64 | — | — | — | 3.25E-02 |  |
| P04906 | Glutathione S-transferase P(GSTP1) | ↓ |  | 0.62 | — | — | 0.50 | 2.41E-02 | tissue<sup>[38]</sup> |
| P13852 | Major prion protein (PRIO) | ↓ | 1 | 0.59 | — | — | — | 3.52E-02 |  |
| P43303 | Interleukin-1 receptor type 2 (IL1R2) | ↓ | 1 | 0.58 | — | — | — | 7.49E-03 | tissue<sup>[39]</sup> |
| B5DFC9 | Nidogen-2 (NID2) | ↓ | 1 | 0.58 | — | — | — | 1.38E-03 | tissue<sup>[40]</sup> |
| Q63556 | Serine protease inhibitor A3M (SPA3M) | ↓ | 1 | 0.58 | — | — | — | 3.48E-04 |  |
| P00786 | Pro-cathepsin H (CATH) | ↓ | 1 | 0.57 | 0.50 | — | — | 5.90E-03 |  |
| P47820 | Angiotensin-converting enzyme (ACE) | ↓ | 1 | 0.55 | — | — | — | 2.73E-02 | serum<sup>[41,42]</sup> |
| Q06000 | Lipoprotein lipase (LIPL) | ↓ | 1 | 0.53 | — | — | — | 1.78E-02 |  |
| O35331 | Pyridoxal kinase (PDXK) | ↓ | 1 | 0.50 | — | — | 0.00 | 2.57E-02 |  |
| Q7TPB4 | CD276 antigen (CD276) | ↓ | 1 | 0.46 | — | — | — | 2.49E-02 |  |
| P14668 | Annexin A5 (ANXA5) | ↓ | 1 | 0.43 | — | — | — | 3.90E-02 |  |
| Q63083 | Nucleobindin-1 (NUCB1) | ↓ | 1 | 0.43 | 0.55 | — | 0.37 | 2.24E-02 |  |
| Q00238 | Intercellular adhesion molecule 1 (ICAM1) | ↓ | 1 | 0.37 | — | — | — | 2.66E-02 | tissue<sup>[43]</sup> |
| Q80YN4 | Atrial natriuretic peptide-converting enzyme (CORIN) | ↓ | 1 | 0.25 | — | — | — | 1.46E-02 |  |
| Q920P6 | Adenosine deaminase (ADA) | ↓ | 1 | 0.00 | — | — | 0.00 | 2.89E-03 | serum, acite<sup>[44]</sup> |
| O08557 | N(G), N(G)-dimethylarginine dimethylamino hydrolase 1 (DDAH1) | ↑ | 1 | — | 7.00 | — | — | 3.27E-02 |  |
| Q8VIF7 | Methanethiol oxidase (SBP1) | ↑ | 1 | — | 4.00 | — | — | 2.13E-02 |  |
| Q63530 | Phosphotriesterase-related protein (PTER) | ↑ | 1 | — | 3.75 | — | — | 1.42E-02 |  |
| P50399 | Rab GDP dissociation inhibitor beta (GDIB) | ↑ | 1 | — | 3.67 | — | — | 1.14E-02 |  |
| Q9WTW7 | Solute carrier family 23 member 1(S23A1) | ↑ | 1 | — | 2.78 | — | — | 3.90E-02 |  |
| Q91XT9 | Neutral ceramidase (ASAH2) | ↑ | 1 | — | 2.50 | — | — | 2.13E-02 |  |
| P09034 | Argininosuccinate synthase (ASSY) | ↑ | 1 | — | 2.00 | — | — | 4.81E-03 | tissue<sup>[45]</sup> |
| Q6AYT0 | Quinone oxidoreductase (QOR) | ↑ | 1 | — | 1.88 | — | — | 3.13E-02 | Cell, tissue<sup>[46]</sup> |
| Q5RLM2 | Solute carrier family 22 member 7 (S22A7) | ↑ | 1 | — | 1.74 | — | — | 4.88E-02 |  |
| Q9JJ40 | Na (+)/H (+) exchange regulatory cofactor (NHRF3) | ↑ | 1 | — | 1.70 | — | — | 3.84E-02 |  |
| P19468 | Glutamate--cysteine ligase catalytic subunit (GSH1) | ↑ | 1 | — | 1.54 | — | — | 2.55E-02 | tissue<sup>[47]</sup> |
| Q64319 | Neutral and basic amino acid transport protein rBAT (SLC31) | ↑ | 1 | — | 1.51 | — | — | 2.22E-02 |  |
| P48199 | C-reactive protein (CRP) | ↓ | 1 | — | 0.66 | — | — | 2.75E-02 | serum<sup>[48]</sup> |
| Q63621 | Interleukin-1 receptor accessory protein (IL1AP) | ↓ | 1 | — | 0.54 | — | — | 3.14E-02 |  |
| P06866 | Haptoglobin chain (HPT) | ↓ | 1 | — | 0.52 | 0.52 | — | 1.57E-02 | serum<sup>[49]</sup> |
| Q9EQV6 | Tripeptidyl-peptidase 1 (TPP-1) | ↓ | 1 | — | 0.50 | — | 0.50 | 2.95E-02 |  |
| Q6IG02 | Keratin(K22E) | ↓ | 1 | — | 0.46 | — | — | 2.49E-02 |  |
| P07171 | Cabining (CALB1) | ↓ | 1 | — | 0.26 | — | 0.00 | 1.32E-02 | tissue<sup>[50]</sup> |
| P26772 | 10 kDa heat shock protein (CH10) | ↓ | 1 | — | 0.17 | — | — | 1.94E-02 |  |
| Q91ZS3 | 45 kDa calcium-binding protein (CAB45) | ↑ | 1 | — | — | 12.0 | — | 3.80E-02 |  |
| Q32KJ6 | N-acetylgalactosamine-6-sulfatase (GALNS) | ↑ | 1 | — | — | 2.14 | — | 3.90E-02 |  |
| P20759 | Ig gamma-1 chain C region (IGHG1) | ↑ | 1 | — | — | 1.86 | 2.77 | 2.52E-02 |  |
| Q6AYS7 | Aminoacylase-1A (ACY1A) | ↑ | 1 | — | — | 1.84 | — | 4.51E-02 |  |
| P48500 | Triosephosphate isomerase (TPIS) | ↑ | 1 | — | — | 1.58 | — | 1.20E-02 | tissue<sup>[51]</sup> |
| Q00657 | Chondroitin sulfate proteoglycan 4 (CSPG4) | ↑ | 1 | — | — | 1.52 | — | 3.36E-02 |  |
| Q63678 | Zinc-alpha-2-glycoprotein (ZA2G) | ↓ | 1 | — | — | 0.64 | — | 2.32E-02 |  |
| P62963 | Profilin-1 (PROF1) | ↓ | 1 | — | — | 0.63 | 0.33 | 3.41E-02 | tissue<sup>[52]</sup> |
| P01026 | Complement C3 (CO3) | ↓ | 1 | — | — | 0.61 | — | 1.05E-02 | tissue<sup>[53]</sup> |
| P20760 | Ig gamma-2A chain C region (IGG2A) | ↑ | 1 | — | — | — | 1.99 | 1.91E-02 |  |
| P22057 | Prostaglandin-H2 D-isomerase (PTGDS) | ↑ | 1 | — | — | — | 1.63 | 1.79E-02 |  |
| P05544 | Serine protease inhibitor A3L(SPA3L) | ↓ | 1 | — | — | — | 0.64 | 9.10E-03 |  |
| P05545 | Serine protease inhibitor A3K (SPA3K) | ↓ | 1 | — | — | — | 0.62 | 2.53E-03 |  |
| P80067 | Dipeptidyl peptidase 1(CATC) | ↓ | 1 | — | — | — | 0.61 | 7.63E-03 |  |
| O70513 | Galectin-3-binding protein (LG3BP) | ↓ | 1 | — | — | — | 0.54 | 4.70E-02 | exosome<sup>[54]</sup> |
| P04797 | Glyceraldehyde-3-phosphate dehydrogenase (G3P) | ↓ | 1 | — | — | — | 0.53 | 2.70E-02 | tissue<sup>[55]</sup> |
| Q9JJ19 | Na (+)/H (+) exchange regulatory cofactor (NHRF1) | ↓ | 1 | — | — | — | 0.51 | 1.17E-02 |  |
| Q02974 | Ketohexokinase (KHK) | ↓ | 1 | — | — | — | 0.50 | 7.76E-03 |  |
| P04642 | L-lactate dehydrogenase A chain (LDH-A) | ↓ | 1 | — | — | — | 0.47 | 1.61E-02 |  |
| O88989 | Malate dehydrogenase, cytoplasmic (MDHC) | ↓ | 1 | — | — | — | 0.45 | 4.53E-02 |  |
| Q99MA2 | Xaa-Pro aminopeptidase 2(XPP2) | ↓ | 1 | — | — | — | 0.43 | 4.23E-02 |  |
| P15473 | Insulin-like growth factor-binding protein 3 (IBP-3) | ↓ | 1 | — | — | — | 0.38 | 4.74E-02 |  |
| Q6AXR4 | Beta-hexosaminidase subunit beta (HEXB) | ↓ | 1 | — | — | — | 0.36 | 2.59E-02 |  |
| Q07936 | Annexin A2 (ANXA2) | ↓ | 1 | — | — | — | 0.33 | 8.05E-03 | tissue<sup>[56]</sup> |
| P24594 | Insulin-like growth factor-binding protein 5 (IBP5) | ↓ | 1 | — | — | — | 0.33 | 3.90E-02 |  |
| P97574 | Stanniocalcin-1 (STC-1) | ↓ | 1 | — | — | — | 0.32 | 1.76E-03 |  |
| Q5XI73 | Rho GDP-dissociation inhibitor 1 (GDIR1) | ↓ | 1 | — | — | — | 0.32 | 3.53E-02 |  |
| Q9R066 | Coxsackievirus and adenovirus receptor homolog (CXAR) | ↓ | 1 | — | — | — | 0.28 | 5.59E-03 |  |
| P04764 | Alpha-enolase (ENOA) | ↓ | 1 | — | — | — | 0.27 | 8.29E-03 | tissue<sup>[51]</sup> |
| P20786 | Platelet-derived growth factor receptor alpha (PGFRA) | ↓ | 1 | — | — | — | 0.27 | 3.29E-02 | tissue<sup>[57]</sup> |
| Q4FZV0 | Beta-mannosidase (MANBA) | ↓ | 1 | — | — | — | 0.26 | 1.14E-02 |  |
| P57097 | Tyrosine-protein kinase (MERTK) | ↓ | 1 | — | — | — | 0.25 | 2.13E-02 |  |
| P19112 | Fructose-1,6-bisphosphatase 1 (F16P1) | ↓ | 1 | — | — | — | 0.21 | 2.85E-02 |  |
| P51635 | Alcohol dehydrogenase [NADP (+)] (AK1A1) | ↓ | 1 | — | — | — | 0.18 | 1.51E-04 |  |
| Q9JLJ3 | 4-trimethylaminobutyaldehyde dehydrogenase (AL9A1) | ↓ | 1 | — | — | — | 0.08 | 3.27E-02 |  |
| Q62638 | Golgi apparatus protein 1 (GSLG1) | ↓ | 1 | — | — | — | 0.07 | 1.52E-05 |  |
| O88767 | Protein/nucleic acid deglycase DJ-1 (PARK7) | ↓ | 1 | — | — | — | 0.07 | 1.14E-02 |  |
| Q812E9 | Neuronal membrane glycoprotein M6-a (GPM6A) | ↓ | 1 | — | — | — | 0.00 | 2.02E-04 |  |

— Indicates that the criteria, compared with day 0, were not met (fold change $\geq 2$ or $\leq 0.5$ , $P < 0.05$ ).

Compared with the profiles before intraperitoneal injection, 19 differential proteins on the 3rd day and 22 differential proteins on 7th day were significantly changed when there was no cancer cell invasion in the pathology of rat abdominal organs, 17 of which were closely associated with ovarian cancer. Four of these have been reported as biomarkers for ovarian cancer. Cell adhesion molecule 4(CADM4) is overexpressed in ovarian tumors, and its cleaved extracellular domain can be detected in the serum of ovarian cancer patients. It is considered to be both a serum biomarker and a therapeutic target for ovarian cancer ^[36]^. Alcohol dehydrogenase 1 (ADH) participates in the metabolism of some biological substances, and its activity is typically increased in ovarian cancer, especially that of the class I isoenzyme, which may be a factor for the metabolic disturbances of important biological substances ^[37]^. It has been reported that intertumor differences in glutathione S-transferase P (GSTP1) expression might influence the response to platinum-based chemotherapy in ovarian cancer patients ^[38]^. Interleukin-1 receptor type 2 (IL1R2) is an important protector against the tumorigenic effects of IL-1 in ovarian cancer, and endometrioid ovarian cancer cells exhibit a specific decrease of IL1R2^[39]^. Nidogen-2 (NID2) is a new promising ovarian malignancy biomarker and is a screening and diagnostic tool for ovarian cancer^[58]^. Serine protease inhibitor A3M (SPA3M) is a member of the serine protease inhibitor superfamily that can regulate cell migration and cell matrix remodeling in cancer, and serine protease inhibitor Kazal type 1 drives proliferation and anoikis resistance in some ovarian cancers ^[59]^. Serum angiotensin-converting enzyme (ACE) levels are increased in patients with epithelial ovarian cancer, and ACE was found to be the key peptidase of the renin-angiotensin system (RAS) that may be associated with ongoing pathobiological events in ovarian carcinogenesis^[42]^. Intercellular adhesion molecule 1 (ICAM1) is increased in serum samples of epithelial ovarian cancer patients and is implicated in tumorigenesis and tumor progression^[43]^. Adenosine deaminase (ADA) levels were found to be higher in the serum and peritoneal fluid of malignant ovarian neoplasm patients, and ADA could be a useful biomarker in the diagnosis and management of ovarian cancer^[44]^. Argininosuccinate synthase (ASSY) is overexpressed in primary ovarian cancer and is regulated by pro-inflammatory cytokines^[45]^, and this trend was also found in our study. Quinone oxidoreductase (QOR) is involved in inflammation and oxidative stress in the development of ovarian cancer^[60]^. It was reported that overexpression of the glutamate-cysteine ligase catalytic subunit (GSH1) protects human ovarian cancer cells against oxidative and gamma-radiation-induced cell death^[47]^. Neutral and basic amino acid transport protein rBAT (SLC31) was reported to be a cysteine transporter and likely plays a major role in the increase of GSH^[61]^. Serum C-reactive protein (CRP) is a widely used biomarker of inflammation and has been previously shown to be a promising biomarker in patients with ovarian cancer^[48]^. Haptoglobin (HPT) is a glycoprotein that regulates the immune response, and serum HPT levels were found to be significantly higher in patients with advanced epithelial ovarian cancer (EOC) with poor survival and were considered a potential biomarker of ovarian carcinoma^[49,62]^. Calbindin (CALB1) is considered to be a mesothelial marker in ovarian cancer^[50]^. The abovementioned urine proteins are closely associated with ovarian cancer and could be used for the detection and prognosis of NuTu-19 cell metastasis progression. This indicates that urine proteomic data could reflect the progression of ovarian cancer.

On the 14th day after intraperitoneal injection, the pathology showed that the omentum was substantially invaded by NuTu-19 cells, and 11 differential proteins were identified in the urine. Among them, three proteins have been reported to be involved in ovarian cancer metastasis and invasion. The expression of triosephosphate isomerase (TPIS) was upregulated in ovarian cancers, and TPIS might represent a biomarker for paclitaxel resistance in ovarian cancer^[51]^. Profìlin-1 (PROF1) is differentially expressed between early- and advanced-stage EOC patient tumor tissues, suggesting a clinical relevance to disease progression, and it is considered as a key regulator of actin cytoskeleton/cell adhesion and cell migration^[52]^. Complement C3 (CO3) is secreted by malignant epithelial cells and promotes EMT in ovarian cancer ^[53]^.

On the 42nd day, the pathology results showed substantial tumor invasion and metastasis in the omentum, ovary, diaphragm, intestine, and liver tissues. A total of 37 differential proteins were identified in the urine, 10 of which have been reported to be involved in the invasion, metastasis and metabolic disorders of ovarian cancer. Galectin-3-binding protein (LG3BP) is enriched in the extracellular vesicles of the ovarian carcinoma and contains sialylated complex N-glycan; it is considered to be a novel potential biomarker for ovarian cancer. Glyceraldehyde-3-phosphate dehydrogenase (G3P) mediates apoptosis in ovarian cancer cells, and high G3P expression levels indicate disease progression in advanced serous ovarian cancer^[55,63]^. These previous reports were consistent with the changes in the urine proteins in the later stage of ovarian cancer in the current study. It is reported that L-lactate dehydrogenase A chain (LDHA) is associated with the induction of ovarian cancer cell apoptosis^[64]^. Malate dehydrogenase (MDHC) is found to be associated with an impaired mitochondrial respiratory capacity in human ovarian and peritoneal cancer cells^[65]^. Annin A2 (ANXA2) is overexpressed in carcinoma tissues compared with normal tissue and has a critical role in EOC cell proliferation; it has the potential to be used as a novel therapeutic target for EOC^[56]^. Insulin-like growth factor-binding protein 5 (IBP5) is overexpressed in high-grade serous carcinomas compared to normal surface epithelium and might play a role in the development of this disease ^[66]^. Alpha-enolase (ENOA) may be involved in the paclitaxel resistance of human ovarian cancer cells, and its expression is higher in metastatic tumors than in primary tumors^[51,67]^. The increasing expression of platelet derived growth factor receptor alpha (PGFRA) is found to be associated with a significantly poorer overall survival of ovarian cancer patients^[57]^. The activity of alcohol dehydrogenase [NADP(+)] (AK1A1) might be an important factor for the disturbances in the metabolism of biological substances in ovarian cancer^[37]^. These urine proteins may indicate a metabolic disorder caused by bloody ascites in the advanced stages (III, IV) of ovarian cancer.

Other differential proteins are found that have not yet been reported to be associated with ovarian cancer but play important roles in the development, progression, migration, invasion and epithelialization of cancer. These proteins may provide important clues regarding the metastasis and invasion of ovarian cancer. For example, the 10 kDa heat shock protein (CH10) identified on days 7 is also known as the cochaperone protein of Hsp60 during the protein folding process, and CH10 is an immune-suppressive growth factor found in maternal serum^[68]^. The upregulated differential protein calcium-binding protein (CAB45) (45 kDa) identified in urine on days 14 has been shown to regulate cancer cell migration through various molecular mechanisms. Moreover, the overexpression of CAB45 results in an altered expression of the molecular mediators of epithelial-mesenchymal transition (EMT), and EMT is an important step in tumor metastasis^[69]^. Prostaglandin-H2D-isomerase (PTGDS) is expressed abundantly in ovarian carcinoma and is regarded as a candidate biomarker for the diagnosis of ovarian carcinoma ^[70]^.

### 3.1.3 Functional analysis of differential urine proteins in rats intraperitoneally injected with NuTu-19 cells

The biological process functional annotation of the differential proteins found on the 7th, 14th, 28th and 42nd day in the intraperitoneal injection group was categorized using DAVID (Fig. 5A); the functional analysis of the differential proteins was investigated using IPA software to explore the biological pathways involved in these differential proteins (Fig. 5B). Seventy-six differential proteins were annotated. Some biological processes were enriched by the differential proteins on the 3rd day. Negative regulation of apoptotic processes and T-cell activation might play roles in cancer cell survival and immunity during ovarian cancer progression. Responses to hypoxia synchronously induce tumor VEGF production, and hypoxia-induced signals would be important factors for initiating and maintaining an active synergistic angiogenic pathway mediated by VEGF in ovarian cancer^[71]^. The biological processes include acute-phase response, positive regulation of nitric oxide biosynthetic process, citrulline metabolic process response to growth hormones and positive regulation of dendrite development, which were all enriched by the differential proteins on the 7th day. Furthermore, the aging process was overrepresented on the 7th and 14th day. The differential proteins involved in biological processes on the 14th day focused on immunoregulation, such as the complement activation classical pathway, the positive regulation of type II hypersensitivity, the innate immune response, the response to magnesium ions, the inflammatory response and the positive regulation of phagocytosis. This indicated that the urine proteins could show a significant immune response on the 14th day after the injection of tumor cells. In 80% of EOC patients, the cancer has usually spread to the omentum by the time of diagnosis^[35]^. In the pathology findings of our study, the omentum was first invaded by on day 14 in the model. The omentum is a lymphoid organ controlling the peritoneal cavity immune response; omental adipocytes promote EOC metastasis by providing energy for rapid tumor growth ^[35,72]^. The pathology changes in ovarian cancer corresponded to the changes in urinary proteins, which suggests that urine could sensitively reflect the immune response process of cancer. The biological processes of the differential proteins on the 42nd day include the carbohydrate metabolic process, the response to hydrogen peroxide, the NAD metabolic process, the positive regulation of the insulin-like growth factor receptor signaling pathway, cellular responses to reactive oxygen species and regulation of the glucose metabolic process. Correspondingly, in the clinical symptoms, the rats in the experimental group on the 42nd day showed cachexia and metabolic disorders, such as anemia, a frog-like abdomen and bloody ascites. The pathological results showed that multiple organs, including the omentum, diaphragm, ovaries, liver and bowl, harbored cancer metastases and invasion. It has been suggested that urinary proteins can reflect the pathological and metabolic disorders of advanced ovarian cancer. The biological processes of negative regulation of cell migration, the cell-cell adhesion regulation of cell shape, and the negative regulation of smooth muscle cell migration are associated with ovarian cancer tumorigenesis, and cell adhesion molecules have been reported to be induced during the early progression of EOC to promote tumor cell migration^[73]^.

**Fig. 5.**
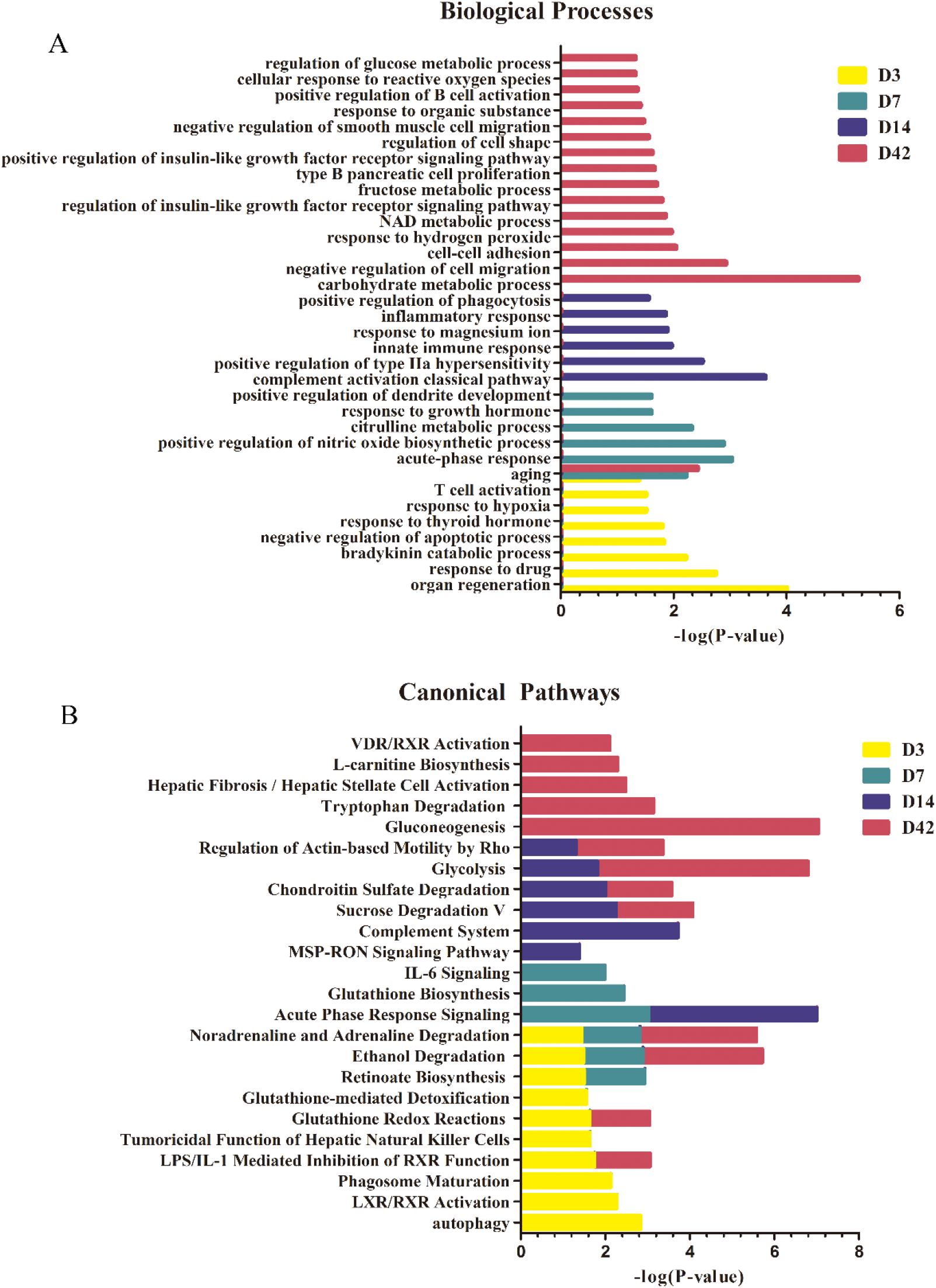
Functional analysis of differential proteins on the 3rd, 7th, 14th and 42nd day in rats intraperitoneally injected with NuTu-19 cells. (A) Biological process; (B) Canonical pathway. (P < 0.05).

Recent evidence has suggested that ovarian cancer cells primary disseminate within the peritoneal cavity and are only superficially invasive^[27]^. Intra-abdominal dissemination occurs at an early stage and is hard to detect. The tumor microenvironment and the involved growth factors and cytokines induced EMT through which tumor cells acquired explosive motility^[26]^. In the canonical pathways of the intraperitoneal injection model, some pathways were closely related to the processes of ovarian cancer, or other cancer, tumorigenesis and invasion.

Among the pathways in which the differential proteins were involved on the 3rd day, it was shown that autophagy could be an effective cancer cell immune escape mechanism, and this mechanism has been implicated in the development of resistance in a variety of cancer types^[74]^. The tumoricidal function of hepatic natural killer cells is related to congenital lymphoid NK cells that have antitumor defense functions, and this pathway mediated antitumor defenses and contributed to the activation and orientation of adaptive immune responses^[75]^. Glutathione redox reactions and the glutathione mediated detoxification pathway were enriched on the 3rd day, and the glutathione biosynthesis pathway was enriched on the 7th day. GSH-activated T cells produce reactive oxygen species (ROS), which triggers an antioxidant glutathione (GSH) response and prevents cell damage; the antioxidant GSH pathway is critical for the inflammatory phase T-cell response by regulating metabolic activity^[76]^. The IL-6 signaling pathway was enriched on the 7th day. Interleukin-6 (IL-6) has been found to be an effective pro-inflammatory cytokine and it plays an important role in the regulation of the immune system in tumorigenesis. According to previous studies, IL-6 transduction is found to mediate tumor cell proliferation and inhibited apoptosis^[77]^. In the MSP-RON signaling pathway (which was enriched on day 28), increasing macrophage stimulating protein (MSP) signaling is elevated in approximately 40% of breast cancers. The activation of the MSP signaling pathway in the bone microenvironment promotes osteolytic bone metastasis^[78]^. The regulated pathway of actin-based movement by Rho was coenriched on the 28th and 42nd days. Recent studies have suggested that the Rho/Rho-associated protein kinases (ROCK) pathway plays a critical role in the regulation of cancer cell motility and invasion, and controlling cell motility via the actin cytoskeleton creates the potential for regulating tumor cell metastasis^[79]^. In the L-carnitine biosynthesis pathway (found on the 42nd day), L-carnitine has a vital role in improving immune system cell function, especially in the lymphocytes, possibly through its antioxidant action^[80]^. In the VDR/RXR activation pathway, RXR is a coactivator of the retinoic acid receptor, and VDR is a vitamin D receptor. The VDR/RXR activation pathway regulates gene expression through a series of metabolic pathways, including the immune response and tumor activation pathways^[81]^.

### 3.2.1 Characterization of orthotopically injected ovarian cancer rat model

After the NuTu-19 cells were orthotopically injected into the left side of the ovary, there were no obvious clinical abnormalities in the experimental group. On the 35th day after injection, the experimental group rats showed rough hair and a reduced diet, but no frog-like abdomen or bloody ascites. Fig. 6 shows the changes in body weight of the o.t.-injected rats. Compared with the body weights of the control rats, there were no significant differences in the body weights of the o.t.-injected experimental group. This indicated that the primary ovarian cancer tumorigenesis was relatively hidden, and any early-stage symptoms were not obvious. Moreover, it also showed that it is not easy to detect ovarian cancer through changes in body weight, and changes in the urine could potentially be seen before there were changes in body weight.

**Fig. 6.**
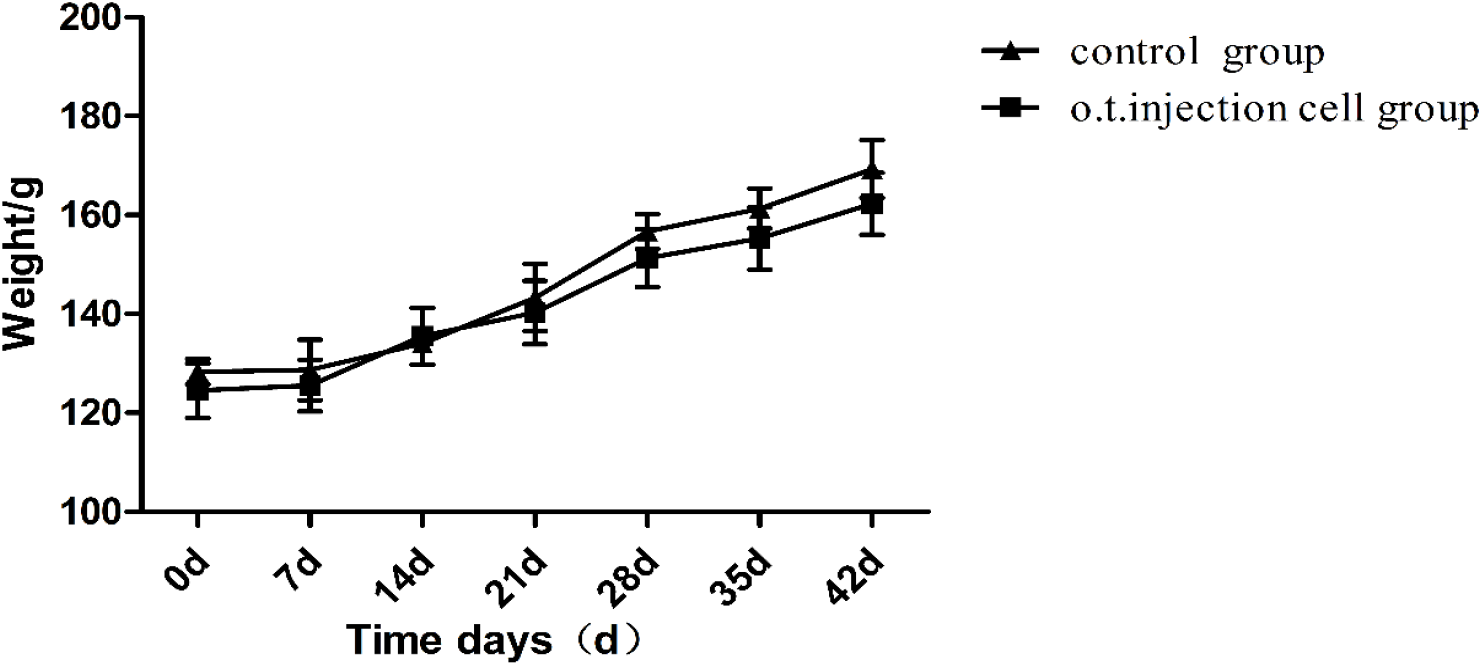
Weight changes in the orthotopically injected ovarian cancer rat model.

The control group presented normal tissues. Histopathological examinations (H&E staining) of the implanted side ovary, unimplanted side ovary, omentum and diaphragm of rats were performed to reveal the progress of the NuTu-19 cells that were orthotopically injected. The control group showed normal tissue results. On day 7 after o.t. injection, the results of the HE staining showed no tumor metastasis of the o.t. group, and tumors were confined to the orthotopic injection side of the ovary compared with the control group (Fig. 7). At 14 days after o.t. injection, there was slight tumor metastasis of the omentum. Twenty-eight days after the injection, the implanted side ovary, unimplanted side ovary (Fig. 7C) and the diaphragm (Fig. 7D) were also invaded by cancer cells. On the 42nd day, the peritoneal organs such as the omentum, diaphragm and ovaries were seriously invaded by tumor cells. Microscopically, the tumor cells were disordered and their size and shape were abnormal. This was consistent with the pathological features of EOC. These pathological changes indicated that the primary ovarian cancer rat model was successfully constructed via the o.t. injection of NuTu-19 cells.

**Fig. 7.**
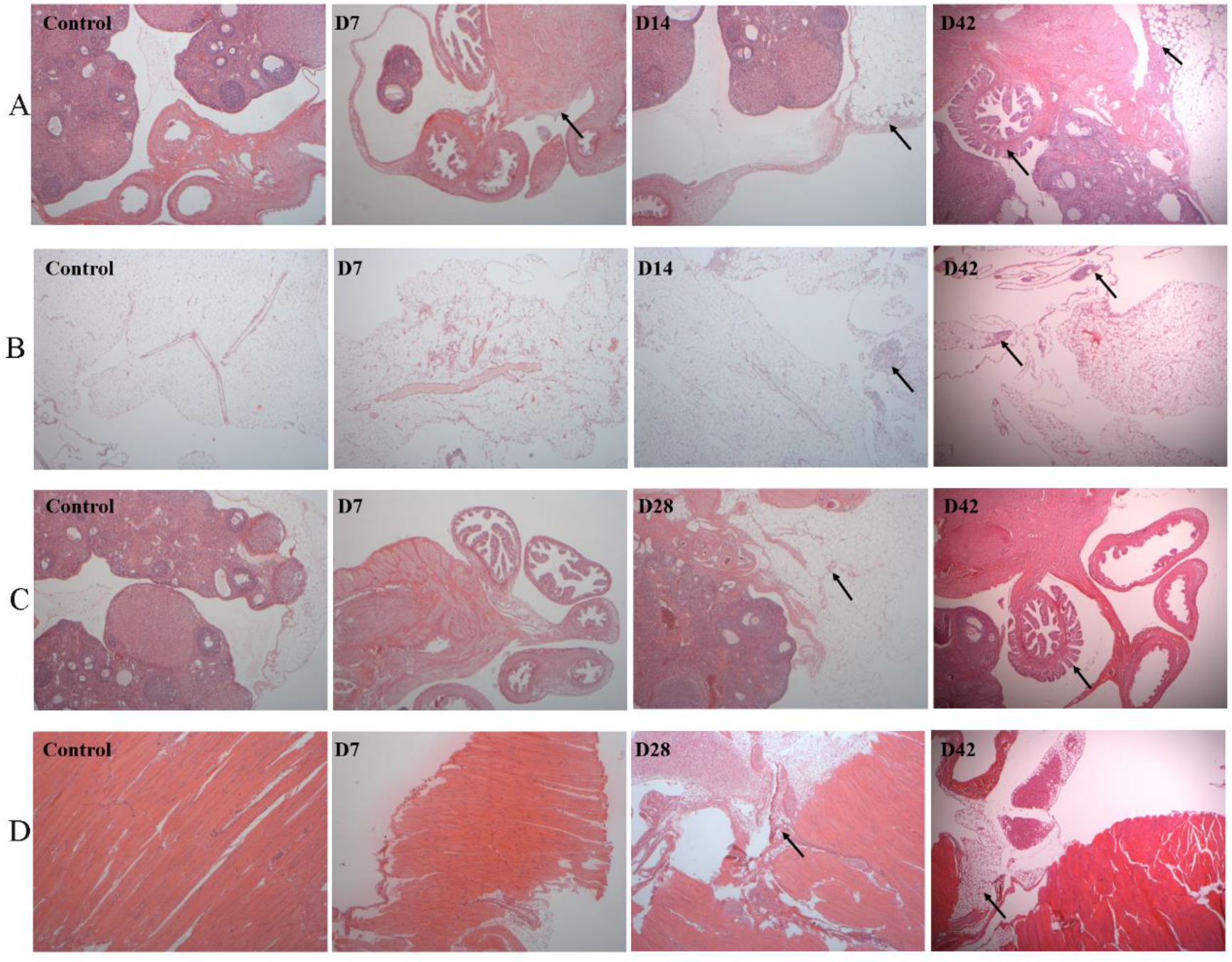
Histopathological characterization of organs in NuTu-19 cell orthotopic injection rat model (40×). Organs were prepared for microscopic examination using H&E stain. Metastases were identified in the A) implanted side ovary, B) omentum, C) unimplanted side ovary and D) diaphragm at different times. Each black arrow denotes the location of the metastasis in the tissue section.

### 3.2.1 Urine proteome changes in the NuTu-19 cell orthotopic injection OC rat model

Twenty urine samples were randomly collected at five time points (on days 7, 14, 28 and 42) from four experimental rats and were analyzed by LC-MS/MS. In total, 602 urinary proteins were identified, and all proteins are listed in Table S3. Among them, 49 differential proteins that had human orthologues were evaluated and had significant changes in all four rats (fold change ≥1.5 or ≤0.67, P < 0.05) (Fig. 8). On the 3rd day, nineteen differential proteins were identified, of which two increased and seventeen of which decreased. On the 7th day, four identified differential proteins were downregulated. On the 14th day, eleven differential proteins were identified, four of which increased and seven of which decreased. On the 28th day, twenty differential proteins were identified, twelve of which increased and eight of which decreased. On the 42nd day, twenty-eight differential proteins were identified, three of which increased and thirty-four of which decreased. The differential proteins are shown in Table S4. The trends of the differential proteins were consistent at each time point in four rats. Interestingly, on the 7th day, no obvious cancer metastases were found in the abdominal organs, and four differential proteins were identified in the urine of these four rats. Among the 49 differential proteins, 24 differential proteins were closely associated with the proliferation and invasion of ovarian cancer, and seven were reported to be biomarkers of ovarian cancer. The urine protein profile changed significantly in the NuTu-19 cell orthotopic injection OC rat model compared to the control.

**Fig. 8.**
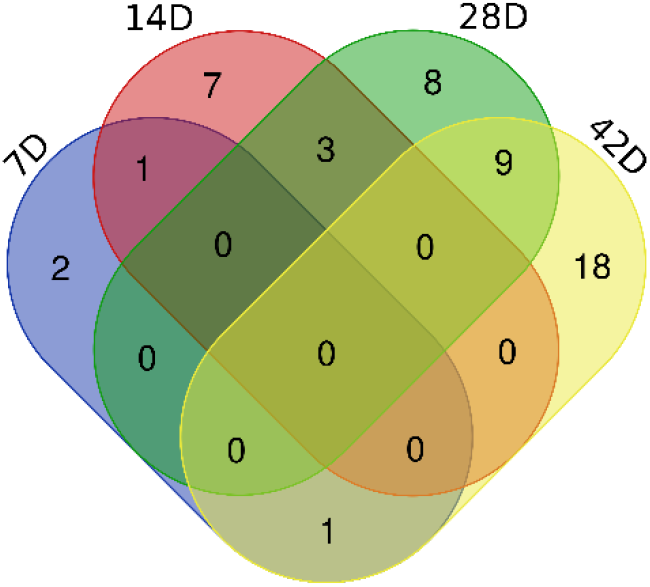
Venn diagram of the differential urinary proteins identified on days 7, 14, 28 and 42 after orthotopic injection of NuTu-19 cells.

**Table 2.**
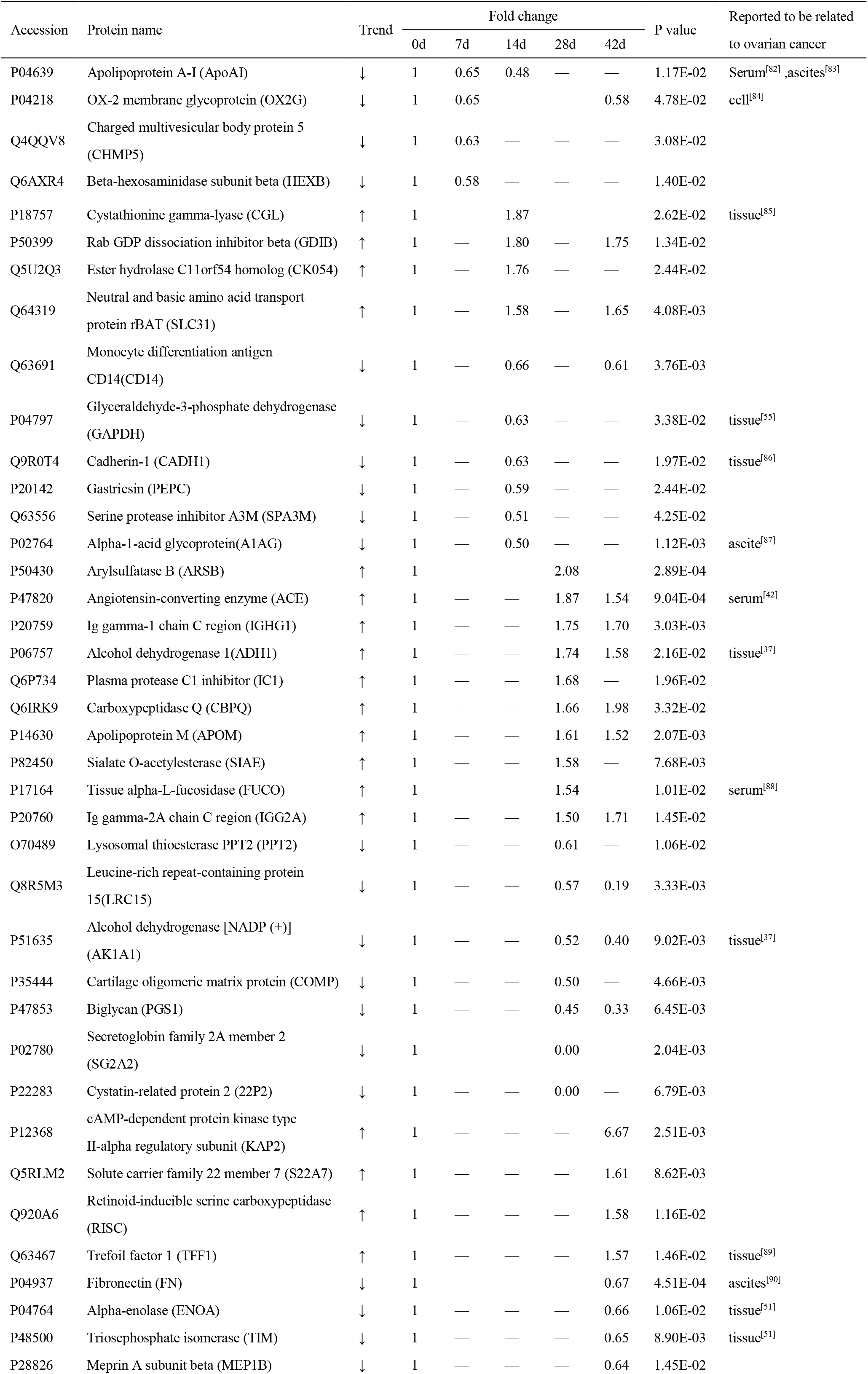
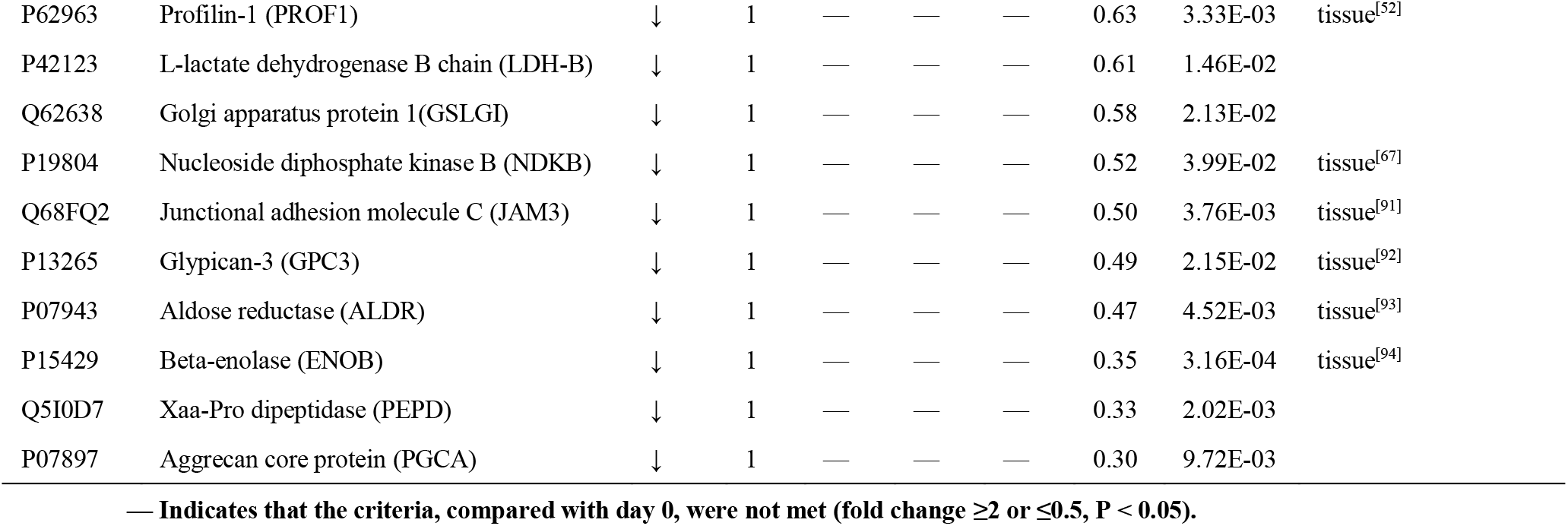
Differential urinary proteins in OC rats orthotopically injected with NuTu-19 cells.

Compared with the self-controls, the urine protein profile of the experimental group significantly changed along with the progression of NuTu-19 cells in the orthotopic injection OC model. On the 7th day after orthotopic injection, there were no tumor metastases in the abdominal organ tissues, and four differential urinary proteins were significantly changed. Among them, two differential proteins were closely related to ovarian cancer. According to recent studies, apolipoprotein A-I (ApoAI) is a high-density lipoprotein that transported excess lipids in the blood to the liver for metabolism. In the current study, ApoAI was found to be decreased in urine. It has been identified as a potential marker for the detection of early-stage serous and endometrioid ovarian cancers^[82]^. The OX-2 membrane glycoprotein (OX2G) is an immunosuppressive factor that is expressed in ovarian tumor cells. The upregulation of OX2G can inhibit the production of Th1 cytokines, which are required for an effective cytotoxic T-cell response. Ovarian tumor cells that express OX2G might potentially suppress antitumor immune responses^[84]^. In addition, charged multivesicular body protein 5 (CHMP5) is essential for the downregulation of receptor signaling during mouse embryo development^[95]^. Beta-hexosaminidase subunit beta (HEXB) helps to regulate the lysosomal β-hexosaminidase α subunit in prostate cancer cells^[96]^. These four differential proteins might be potential biomarkers for the early diagnosis of ovarian cancer.

Eleven homologous differential proteins were identified on the 14th day, seven of which were differentially expressed in the blood or tissue samples of the orthotopic injection OC model. In addition to these, three differential proteins were also identified in the intraperitoneal injection model: the neutral and basic amino acid transporter rBAT (SLC31), glyceraldehyde-3-phosphate dehydrogenase (G3P), and serine protease inhibitor A3M (SPA3M). Cystathionine γ-lyase (CGL) is an enzyme that is involved in the synthesis of cysteine and has been reported to be a marker for clear cell carcinoma in a proteomic screen; CGL was found to be expressed at high levels in clear cell carcinomas of the ovary and endometrium ^[85]^. Monocyte differentiation antigen (CD14) cells were reportedly found in ascites of ovarian tumors, and CD14 significantly inhibits the antigen-specific CD4(+) T-cell immune response^[97]^. CD14 also regulates cancer cell EMT and invasion in vitro^[98]^. Glyceraldehyde-3-phosphate dehydrogenase (GAPDH) is reportedly involved in cellular metabolism, and the overexpression of GAPDH increases ovarian cancer cell apoptosis. This enzyme is considered to be a metabolic marker in advanced serous ovarian cancer^[55,63]^. Cadherin 1 (CADH1) has been reported to have downregulated expression in ovarian cancer tissues and participates in the epithelial-mesenchymal transition; P-cadherin had been reported to induce the early progression of EOC and to promote tumor cell migration^[99]^. The alpha-1-acid glycoprotein (A1AG) is reportedly an immunosuppressive protein found in the ascites of ovarian cancer and inhibits the secretion of IL-2 by lymphocytes to achieve immunosuppressive effects^[87]^. A1AG is used as a serum biomarker in lung cancer ^[100]^.

Twenty differential proteins were identified on the 28th day, five of which were explicitly reported to be associated with ovarian cancer. Angiotensin-converting enzyme (ACE) is a key enzyme in the renin-angiotensin system (RAS) of the female reproductive system and evidence implicates ACE in the pathophysiology of carcinogenesis^[101]^. ACE is a reported marker for disseminated germinoma tumors and was found to be useful for monitoring treatment in ovarian cancer^[41]^. High expression of alcohol dehydrogenase 1 (ADH) is found to be significantly associated with the risk of advanced serous ovarian cancer^[102]^. It has been reported that the alcohol dehydrogenases, especially the ADH class I isoenzymes, have increased activity in ovarian cancer, and alcohol dehydrogenase [NADP(+)] (AK1A1) and ADH are related to metabolic disorders^[37]^. Plasma protease C1 inhibitor (IC1) is enhanced and released by drug-resistant ovarian cancer cells^[103]^. In addition, IC1 is a molecular indicator used for screening early breast cancer^[104]^. The activity of α-L-fucosidase (FUCO) is found to be increased in ovarian cancer tissues, and FUCO plays a vital role in inhibiting macrophage migration^[88]^. Biglycan (PGS1) is overexpressed in ovarian cancer tissue and is a specific marker and an autocrine angiogenic factor of tumor endothelial cells^[105]^.

On the 42nd day, twenty-eight differential proteins were identified, 10 of which were previously reported to be associated with ovarian cancer. Three of these differential proteins, including α-enolase (ENOA), triose phosphate isomerase (TPIS), and prefibrin-1 (PROF1), were identified in the i.p. injection OC model. Trefoil factor 1 (TFF1) is a secreted protein that is highly expressed in mucinous ovarian cancer and enhances the malignant phenotype of mucinous ovarian cancer cells through Wnt/β-catenin signaling ^[89]^. Fibronectin (FINC) has been reported to promote early ovarian cancer metastasis by upregulating FAK-PI3K/Akt migration and invasion of ovarian cancer cells^[106]^ and is considered as an ovarian cancer biomarker for monitoring and recurrence^[107]^. Nucleoside diphosphate kinase B (NDKB) is a nuclear nucleoside diphosphate kinase subunit. It affects cell development and proliferation and is associated with the paclitaxel-resistance of human ovarian cancer cells^[67]^. Cross-linked adhesion molecule C (JAMC) is a transmembrane protein that plays an important role in regulating immune cell recruitment and angiogenesis in endothelial cells. JAMC is closely related to tumor growth and invasiveness^[91]^. Phosphatidylinositol-3 (GPC3) is involved in cell proliferation, adhesion, migration, invasion and differentiation during the development of most mesoderm tissues and organs. The downregulation of phosphatidylinositol-3 expression increased the migration, invasion and tumorigenicity of human ovarian cancer cells^[92]^. Beta-enolase (ENOB) has muscle specificity and is regarded as a biomarker expressed in nonhuman primate ovarian sex cord-stromal tumors^[94]^. In addition, ENOB is a tumor marker for the diagnosis and prognosis of small cell lung cancer^[108]^.

Additionally, some other differential proteins might be potential novel ovarian cancer markers. For example, the leucine-rich repeat protein 15 (LRC15) identified on the 28th day was reportedly downregulated and found to be correlated with tumor invasion and metastasis ^[109]^. Meprin (MEP1B) was identified on day 42, and this protein is a membrane matrix metalloproteinase expressed in epithelial tissues and cancer cells and could play a role in epithelial differentiation and cell migration^[110,111]^.

### 3.2.3 Functional analysis of differential urine proteins in rats orthotopically injected with NuTu-19 cells

The differential proteins identified on the 7th, 14th, 28th and 42nd days were analyzed for their biological processes by the DAVID database (Fig. 9A), and pathway analysis was performed using IPA software (Fig. 9B). Regarding the biological processes, the differential proteins found on the 7th day were involved in lipid storage and lysosomal tissue processes. Adipocytes provide fatty acids and other substances for the rapid growth of tumors, and adipocytes are important components in the tumor microenvironment and contribute to the metastasis of cancer cells to the omentum^[26]^. The differential proteins on the 14th day were involved in drug response, organ regeneration, and carbohydrate metabolism. On the 28th day, the biological processes were related to the immune system and immune regulation, as well as the classical pathways of complement activation, positive regulation of B-cell activation, peptide catabolism, phagocytosis and negative regulation of the JAK-STAT cascade. The negative regulation of the JAK-STAT cascade is enriched, and Jak/STAT signaling is involved in various types of blood cell disorders and cancers in humans; the activation of Jak/STAT signaling is associated with more invasive metastatic cancers^[112]^. The JAK-mediated signaling protein cascade activates cytokines and their receptors, and STAT1 is involved in the regulation of cell growth, apoptosis and differentiation; the inactivation and mutation of its negative regulatory molecules causes dysfunction^[113]^. On the 42nd day, the enriched biological processes were related to metabolism, including glycolysis, cellular response to glucose stimulus, retinoic acid metabolism, response to thyroid hormone, positive regulation of B-cell activation and peptide catabolic processes.

**Fig. 9.**
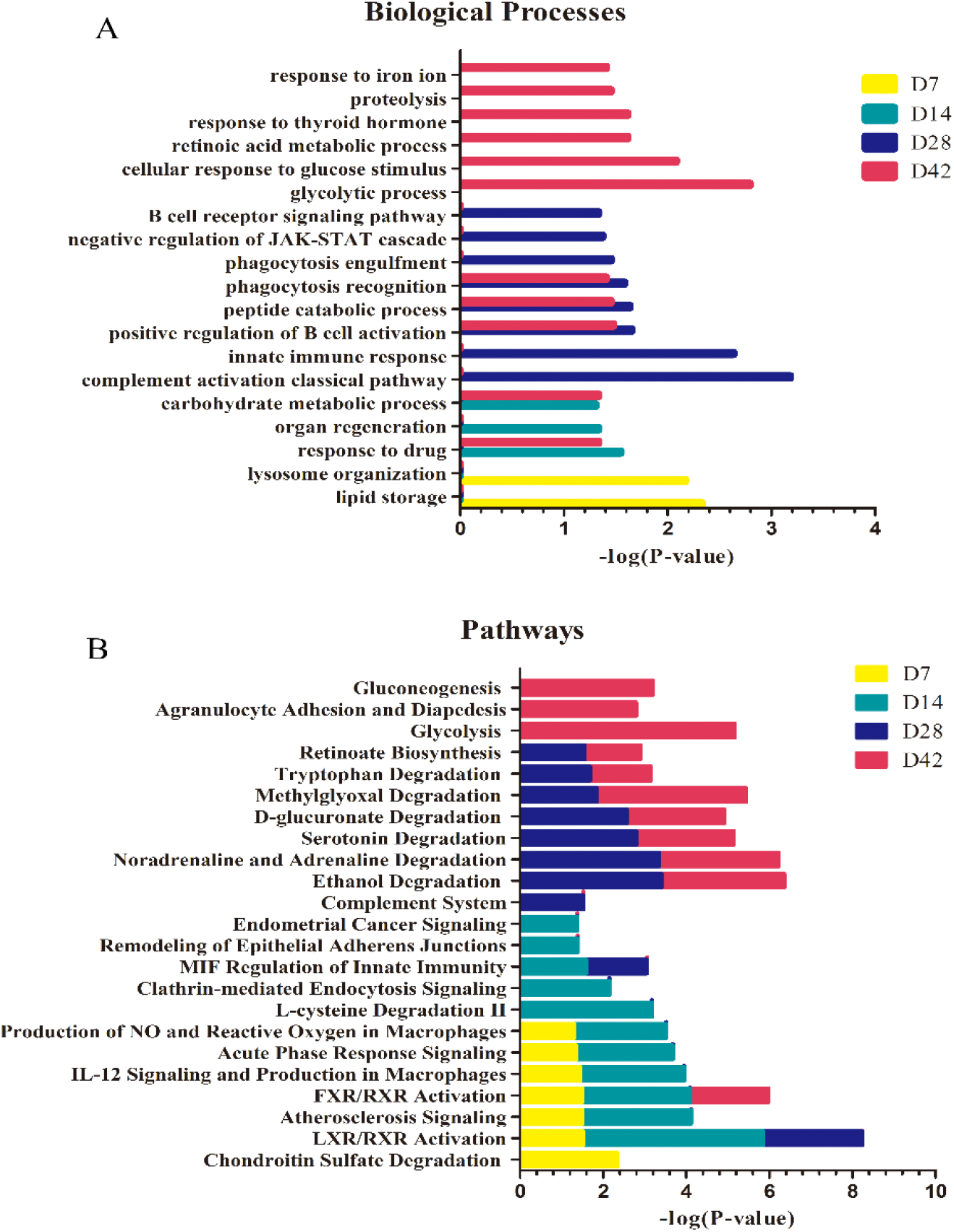
Functional analysis of differential proteins on days 7, 14, 28 and 42 in rats orthotopically injected with NuTu-19 cells. (A) Biological processes; (B) Canonical pathways. (P < 0.05).

Primary ovarian cancer metastasizes via a passive mechanism by which cancer cells shed from the primary tumor and are carried by the physiological movement of peritoneal fluid to the omentum. Hematogenous metastasis of ovarian cancer to the omentum occurs via circulating tumor cells. Subsequently, the cancer cells exit the metastatic site by extravasation and establish a metastatic tumor nodule ^[114]^.

The classical pathways that are significantly associated with all of the differential proteins are shown (Fig. 9B). On the 7th and 14th days, FXR/RXR (the farnesoid X receptor/retinoid X receptor-like) pathway that regulates lipoprotein and lipid metabolism was commonly enriched ^[115]^. Adipocytes in the membrane were shown to promote the proliferation and migration of ovarian cancer cells, which preferentially invaded the omentum^[116]^. Some reports have indicated that cancer cells induce metabolic changes in adipocytes and stimulate the defatting of mature differentiated adipocytes to release lipids to promote tumor progression^[117]^. IL-12 is a glycoprotein produced by macrophages and B lymphocytes. The IL-12 cytokine family are crucial factors that regulate the T-cell response^[118]^. Macrophages are stimulated by immune factors and inflammation secrete of a variety of biologically active substances, such as NO and ROS. NO is an important immunomodulatory factor that is closely related to tumor development ^[119]^. The L-cysteine degradation pathway is a key mechanism for affecting tumor cells, and this pathway has the ability to regulate tumor cells to detoxify ROS ^[120]^.

The regulatory pathway for innate immunity was enriched on the 14th and 28th days. The generation of spontaneous T-cell responses able to defeat tumor-associated antigens depends on innate immune activation. According to a recent study, macrophage migration inhibitory factor (MIF) was recognized as playing an important role in both innate and acquired immunity. The research showed that MIF could also stimulate tumor cell proliferation and differentiation and inhibited tumor cell apoptosis; MIF was also found to play a key role in cell cycle regulation and tumorigenesis ^[121]^.

The remodeling pathway of epithelial adhesions was enriched on the 28th day, and this pathway plays a key role in the remodeling of neoplastic epithelium during metastasis and regulates EOC growth and transmission^[122]^. Abnormal epithelial differentiation is an early event in ovarian cancer^[73]^. In the serotonin degradation pathway, serotonin typically exhibits growth stimulatory effects in invasive cancers through the 5-HT1 and 5-HT2 receptor; in contrast, low doses of serotonin could inhibit tumor growth by reducing the blood supply to the tumor^[123]^.

The granulocyte adhesion and permeation pathways were enriched on the 42nd day. It has been reported that integrin-mediated strong adhesion could induce rapid granulocyte adhesion and promoted leukocyte adhesion to the inner wall of blood vessels, which was associated with granulocyte entry into the interstitial area, injury and immune responses to stress sites^[124]^.

### 3.3 Comparison of intraperitoneal injection and orthotopic injection of NuTu-19 cells in two ovarian cancer rat models

According to the pathological processes (Fig. 10), the disease progression of the two models was different. As shown in Fig. 11, the urinary protein changes in the intraperitoneal injection and orthotopic injection OC models were different, and the overlap ratio of proteins was low. The repeated proteins included SLC31, ADH1, TPIS, IGHG1, IGG2A, AK1A1, S22A7, ENOA, ACE, IC1, PROF1, HEXB, G3P, GDIB, SPA3M, and GSLG1. Unexpectedly, overlapping proteins were identified at interlaced time points in the two models, and only six differential proteins (PROF1, IGHG1, IGG2A, AK1A1, GSLG1, and ENOA) were identified on the 42nd day. The possible reason for this is, on the one hand, the same tumor cells were injected at different positions to cause urinary protein changes and these changes were related to the ovary with complex tissue types. On the other hand, the different number of cancer cells injected may have induced different disease progressions. Tumor growth depends not only on the genetic changes in malignant cells but also on changes in the tumor microenvironment (TME), such as the matrix, blood vessels, infiltrating inflammatory cells, chronic inflammation and evasion of antitumor immune response, which is necessary for tumorigenesis and cancer progression; therefore, immunity and inflammation are the two cores that constitute the tumor microenvironment^[125]^. Interestingly, the functional annotations of the differential proteins in the two models were different, i.e., most of the involved biological processes and pathways were different, but some common biological pathways were also identified. For example, in the chondroitin sulfate degradation pathway, chondroitin sulfate (CS) is a type of sulfated polysaccharide, and CS is abundantly present in the extracellular matrix of ovarian cancer. It had been demonstrated that structural changes in CS play a crucial role in cancer development and progression and regulate cell adhesion, proliferation, migration and tumor blood vessels^[126]^. In the LXR/RXR activation pathway, LXR/RXR is involved in lipid metabolism, including inflammatory responses and cholesterol metabolism^[115]^. Studies of the comparative proteomic analysis of urine revealed a downregulation of acute-phase response signals and LXR/RXR activation pathways in prostate cancer^[127]^. Some of the identified pathways were involved in the body’s immune response. For example, the acute-phase response signal pathway (involving fibrinogen, alpha-1-acid glycoprotein and haptoglobin) plays an important role in nonspecific immunity during tumorigenesis^[113]^. In the classical pathway of complement activation (enriched in the orthotopic injection model on the 28th day and in the intraperitoneal injection model on the 14th day), this pathway often plays a role in endometriosis-like ovarian cancer^[128]^. It is involved in mechanisms effected by the innate immune system, and cancer cells secrete complement proteins, so complement likely plays an important role in stimulating tumor cell proliferation to enhance cancer growth ^[129]^. Complement activation is also involved in adaptive immune responses, and complement activation in the tumor microenvironment enhances tumor growth and metastasis^[130]^. The activation of the innate immune response determines the production of a spontaneous T-cell response against tumor-associated antigens and kills homologous tumor cells ^[131]^. The innate immune system not only plays an important part in tumor promotion but also has a key role in the development of antitumor adaptive immune responses^[125]^. In the positive regulation of the B-cell activation pathway, some activated B cells produce a large number of cytokines, and B cells participate in antitumor effects such as immune regulation and facilitate the development and maintenance of regulatory T cells^[132]^. Some biological processes were commonly enriched in two models, including the response to drugs, organ regeneration, carbohydrate metabolism, innate immune response, positive regulation of B-cell activation, and the response to thyroid hormone.

**Fig. 10.**
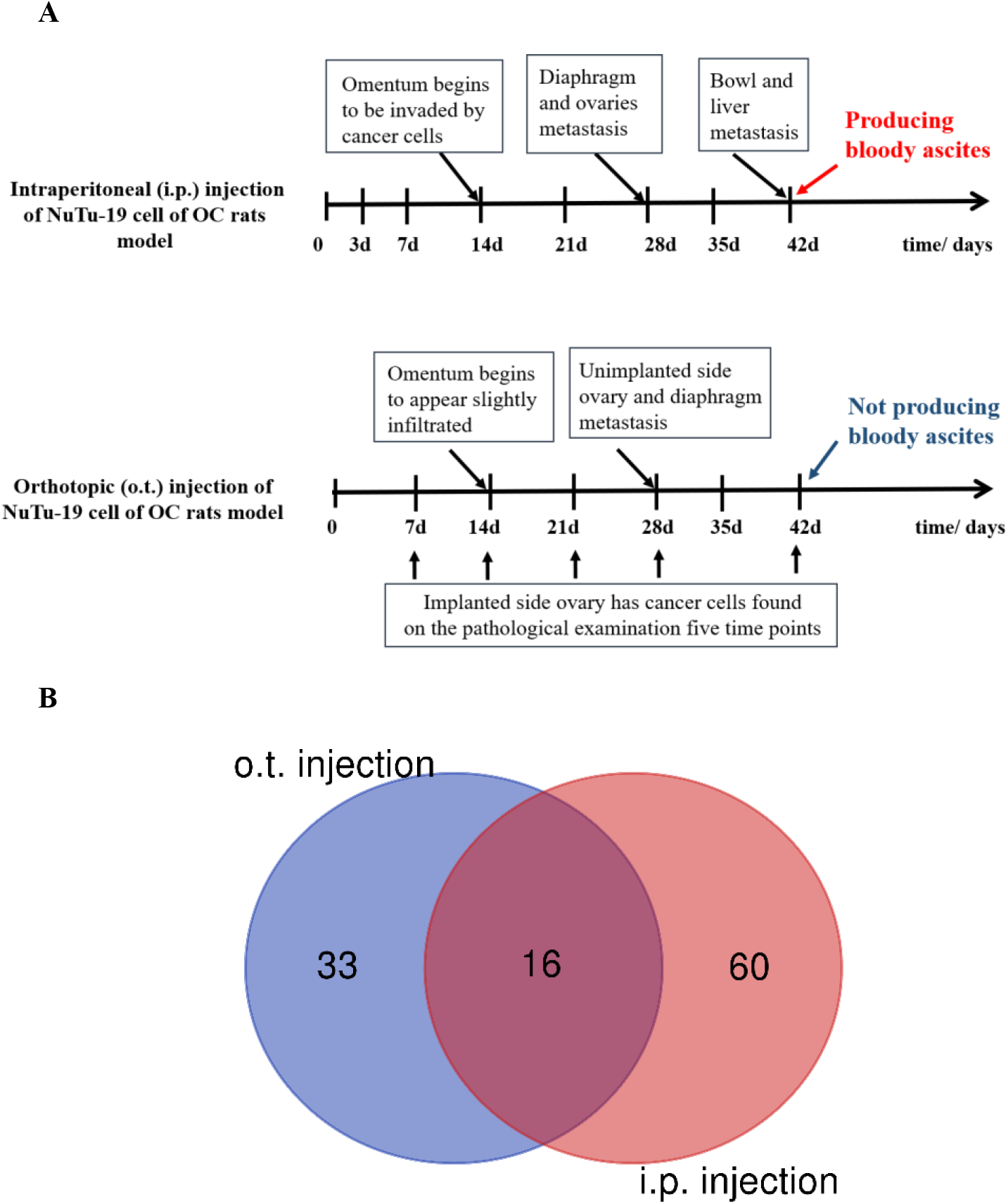
Comparison of pathological processes and urinary differential proteins between the intraperitoneal (i.p.) injection OC model and orthotopic (o.t.) injection OC model. A) Comparison of pathological disease progression in the two models; B) Venn diagram of differential urinary proteins in the two models.

**Fig. 11.**
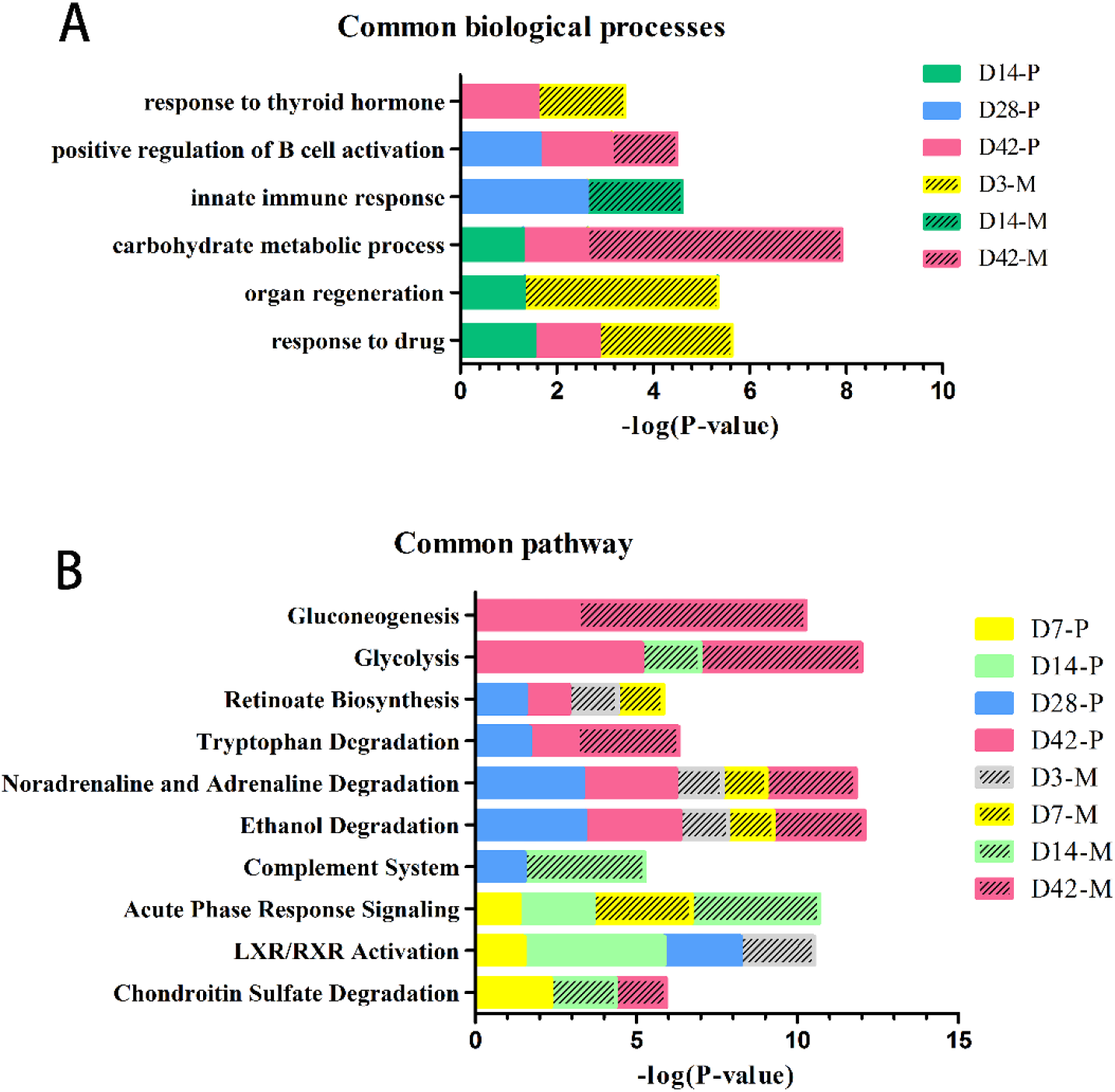
Comparison of common biological processes and classical pathways of differential urinary proteins between the intraperitoneal (i.p.) injection OC model and the orthotopic (o.t.) injection OC model. A) Common biological process; B) Common pathways. M indicates the intraperitoneal injection model and P indicates the orthotopic injection model. D7-P indicates the orthotopic injection model on day 7; D7-M indicates the intraperitoneal injection model on day 7. P <0.05.

Interestingly, the differential urinary proteins on the 14th day of the intraperitoneal injection model were involved with reactions and pathways such as immune regulation and complement activation. However, the orthotopic injection ovarian cancer model on the 28th day was related to the immune system and immune regulation. Biological processes and pathways such as immunoregulation, antibody-antigen binding and the positive regulation of B-cell activation indicated that the disease progression was different in two ovarian cancer models.

Other common pathways were involved in tumor metabolism; for example, in the norepinephrine and adrenaline degradation pathways. Studies have shown that adrenaline and norepinephrine protect ovarian cancer cells from anoikis^[133]^. Regarding the tryptophan degradation pathway, the tryptophan-degrading enzyme guanamine 2,3-dioxygenase (IDO) participates in an antimicrobial and antitumor immune effect mechanism, and accelerated tryptophan degradation seems to be represent an immune escape mechanism^[134]^. Notably, the two models were enriched in the glycolytic pathway on the 42th day. The cancer cells strongly upregulated glucose uptake and glycolysis, which resulted in an increased production of the intermediate glycolysis metabolites and the final product pyruvate. Glycolysis intermediates provide the necessary anabolism for cell proliferation and tumor growth^[135]^. It has been reported that the gluconeogenesis pathway of monocyte anabolism is involved in the inflammatory tumor microenvironment and is a key event in the development of tumors. The gluconeogenesis pathway accelerated tumor growth and suggests that metabolic disorders are associated with advanced cancer^[136]^. Surprisingly, the differential proteins on the 42nd day in two models were involved in a variety of biological processes and classical pathways of glucose metabolism and amino acid metabolism. This indicated that the urine could reflect the progression of the metabolic disorders of advanced ovarian cancer.

Interestingly, some differential urinary proteins found in this study have not been reported in previous studies. Urine can reflect changes in the body, and the use of an animal model excludes the interference of confounding factors. The previously unreported proteins, identified here with good credibility, might be new urinary protein biomarkers of ovarian cancer. Biological repeat experiments were added to the study, and this study lays the foundation for their future clinical validation. Although the intra-abdominal dissemination of ovarian cancer cells had similar clinical manifestations after the metastasis of the primary ovarian cancer, the early treatment methods and prognostic methods are quite different. This study might support the differential diagnosis of primary ovarian cancer and ovarian peritoneal carcinomatosis.

The successful clinical management of patients with epithelial ovarian cancer is limited by the lack of a reliable and specific method for early detection and the frequent recurrence of chemoresistant disease^[15]^. Alternative biomarker combinations might lead to significant improvements in the detection of ovarian cancer, and we have reason to expect that a panel of differential urinary protein combinations may provide earlier and more sensitive clues regarding the physiological progression and early clinical diagnosis of ovarian cancer^[137]^. There was a diverse pathogenesis of the disease at different time points, which showed different combinations of urine proteins. In particular, in the orthotopic injection model of NuTu-19 cells, the differential urinary proteins provided diagnostic clues for ovarian cancer on days 7, 14, 28, and 42, while the pathology offered clues on the 14th day; the urine proteins on days 7 and 14 could supply valuable information for an earlier diagnosis. A prospective clinical analysis of the panel of urinary protein is needed to confirm this strategy as an effective screening tool for early-stage ovarian cancer. The development of mass spectrometry technology has made it possible to rapidly and conveniently screen urine samples for diseases. A simple and novel membrane storage technique of urine preservation could be an optimal choice for storing regularly collected human urine samples^[138]^. This makes it possible to conveniently store a large number of urine samples from various time points to form a personalized urine sample profile, which provides the possibility for the early detection and early treatment of diseases.

## 4. Conclusion

This study demonstrates that urinary proteins can sensitively reflect early changes in an orthotopic ovarian cancer rat model. The urinary proteins could reflect the progression of ovarian cancer in both the intraperitoneal and orthotopic models. Importantly, in the orthotopic model, differential proteins APOA1, OX2G, CHMP5, and HEXB were identified in the urine before ovarian cancer metastasis.

## Author contributions

Y.L and Y.G. conceived and designed the experiments. Y.L, L.Z. and W.M. performed the experiments. Y.L analyzed the data and wrote the manuscript. All authors approved the final manuscript.

## Acknowledgments

This work was supported by the National Key Research and Development Program of China (2018YFC0910202, 2016YFC1306300), Beijing Natural Science Foundation (7172076), Beijing cooperative construction project (110651103, Beijing Normal University (11100704), and Peking Union Medical College Hospital (2016-2.27).

## Reference

[1] Siegel R L, Miller K D, Jemal A. Cancer Statistics, 2017[J]. CA Cancer J Clin, 2017, 67(1): 7–30.

[2] Dong X, Men X, Zhang W, et al. Advances in tumor markers of ovarian cancer for early diagnosis[J]. Indian J Cancer, 2014, 51 Suppl 3: e72–6.

[3] Gloss B S, Samimi G. Epigenetic biomarkers in epithelial ovarian cancer[J]. Cancer Lett, 2014, 342(2): 257–63.

[4] Argento M1 H P, Gauchez As. Ovarian cancer detection and treatment: current situation and future prospects.[J]. Anticancer Research, 2008, 28: 3135–3138.

[5] Menon U, Griffin M, Gentry-Maharaj A. Ovarian cancer screening--current status, future directions[J]. Gynecol Oncol, 2014, 132(2): 490–5.

[6] Doubeni C A, Doubeni A R, Myers A E. Diagnosis and Management of Ovarian Cancer[J]. Am Fam Physician, 2016, 93(11): 937–44.

[7] Gao Y. Urine-an untapped goldmine for biomarker discovery?[J]. Sci China Life Sci, 2013, 56(12): 1145–6.

[8] Zhao M, Li M, Yang Y, et al. A comprehensive analysis and annotation of human normal urinary proteome[J]. Sci Rep, 2017, 7(1): 3024.

[9] Badgwell D, Lu Z, Cole L, et al. Urinary mesothelin provides greater sensitivity for early stage ovarian cancer than serum mesothelin, urinary hCG free beta subunit and urinary hCG beta core fragment[J]. Gynecol Oncol, 2007, 106(3): 490–7.

[10] Wu J, Gao Y. Physiological conditions can be reflected in human urine proteome and metabolome[J]. Expert Rev Proteomics, 2015, 12(6): 623–36.

[11] Liao J B, Yip Y Y, Swisher E M, et al. Detection of the HE4 protein in urine as a biomarker for ovarian neoplasms: Clinical correlates[J]. Gynecol Oncol, 2015, 137(3): 430–5.

[12] Ye B, Skates S, Mok S C, et al. Proteomic-based discovery and characterization of glycosylated eosinophil-derived neurotoxin and COOH-terminal osteopontin fragments for ovarian cancer in urine[J]. Clin Cancer Res, 2006, 12(2): 432–41.

[13] Anderson N S, Bermudez Y, Badgwell D, et al. Urinary levels of Bcl-2 are elevated in ovarian cancer patients[J]. Gynecol Oncol, 2009, 112(1): 60–7.

[14] Zhao M. Dynamic changes of urinary proteins in focal segmental glomerulosclerosis model[J]. Adv Exp Med Biol, 2015, 845: 167–73.

[15] Garson K, Shaw T J, Clark K V, et al. Models of ovarian cancer--are we there yet?[J]. Mol Cell Endocrinol, 2005, 239(1-2): 15–26.

[16] Yuan Y, Zhang F, Wu J, et al. Urinary candidate biomarker discovery in a rat unilateral ureteral obstruction model[J]. Sci Rep, 2015, 5: 9314.

[17] Wu J, Li X, Zhao M, et al. Early Detection of Urinary Proteome Biomarkers for Effective Early Treatment of Pulmonary Fibrosis in a Rat Model[J]. Proteomics Clin Appl, 2017, 11(11-12).

[18] Zhang F, Ni Y, Yuan Y, et al. Early urinary candidate biomarker discovery in a rat thioacetamide-induced liver fibrosis model[J]. Sci China Life Sci, 2018, 61(11): 1369–1381.

[19] Zhao M, Wu J, Li X, et al. Urinary candidate biomarkers in an experimental autoimmune myocarditis rat model[J]. J Proteomics, 2018, 179: 71–79.

[20] Wang Y, Chen Y, Zhang Y, et al. Differential ConA-enriched urinary proteome in rat experimental glomerular diseases[J]. Biochem Biophys Res Commun, 2008, 371(3): 385–90.

[21] Zhang L, Li Y, Gao Y. Early changes in the urine proteome in a diethyldithiocarbamate-induced chronic pancreatitis rat model[J]. J Proteomics, 2018, 186: 8–14.

[22] Zhang F, Wei J, Li X, et al. Early Candidate Urine Biomarkers for Detecting Alzheimer’s Disease Before Amyloid-beta Plaque Deposition in an APP (swe)/PSEN1dE9 Transgenic Mouse Model[J]. J Alzheimers Dis, 2018, 66(2): 613–637.

[23] Wu J, Guo Z, Gao Y. Dynamic changes of urine proteome in a Walker 256 tumor-bearing rat model[J]. Cancer Med, 2017, 6(11): 2713–2722.

[24] Ramalingam P. Morphologic, Immunophenotypic, and Molecular Features of Epithelial Ovarian Cancer[J]. Oncology (Williston Park), 2016, 30(2): 166–76.

[25] Rose G S, Tocco L M, Granger G A, et al. Development and characterization of a clinically useful animal model of epithelial ovarian cancer in the Fischer 344 rat[J]. Am J Obstet Gynecol, 1996, 175(3 Pt 1): 593–9.

[26] Nakayama K, Nakayama N, Katagiri H, et al. Mechanisms of ovarian cancer metastasis: biochemical pathways[J]. Int J Mol Sci, 2012, 13(9): 11705–17.

[27] Lengyel E. Ovarian cancer development and metastasis[J]. Am J Pathol, 2010, 177(3): 1053–64.

[28] Azais H, Queniat G, Bonner C, et al. Fischer 344 Rat: A Preclinical Model for Epithelial Ovarian Cancer Folate-Targeted Therapy[J]. Int J Gynecol Cancer, 2015, 25(7): 1194–200.

[29] Fan L, Liu Y, Zhang X, et al. Establishment of Fischer 344 rat model of ovarian cancer with lymphatic metastasis[J]. Arch Gynecol Obstet, 2014, 289(1): 149–54.

[30] Sloan Stakleff K D, Rouse A G, Ryan A P, et al. A novel early-stage orthotopic model for ovarian cancer in the Fischer 344 rat[J]. Int J Gynecol Cancer, 2005, 15(2): 246–54.

[31] Wisniewski J R, Zougman A, Nagaraj N, et al. Universal sample preparation method for proteome analysis[J]. Nat Methods, 2009, 6(5): 359–62.

[32] Sun W, Li F, Wu S, et al. Human urine proteome analysis by three separation approaches[J]. Proteomics, 2005, 5(18): 4994–5001.

[33] Old W M, Meyer-Arendt K, Aveline-Wolf L, et al. Comparison of label-free methods for quantifying human proteins by shotgun proteomics[J]. Mol Cell Proteomics, 2005, 4(10): 1487–502.

[34] Schmidt C, Gronborg M, Deckert J, et al. Mass spectrometry-based relative quantification of proteins in precatalytic and catalytically active spliceosomes by metabolic labeling (SILAC), chemical labeling (iTRAQ), and label-free spectral count[J]. RNA, 2014, 20(3): 406–20.

[35] Lungchukiet P, Sun Y, Kasiappan R, et al. Suppression of epithelial ovarian cancer invasion into the omentum by 1alpha,25-dihydroxyvitamin D3 and its receptor[J]. J Steroid Biochem Mol Biol, 2015, 148: 138–47.

[36] Buchanan P C, Boylan K L M, Walcheck B, et al. Ectodomain shedding of the cell adhesion molecule Nectin-4 in ovarian cancer is mediated by ADAM10 and ADAM17[J]. J Biol Chem, 2017, 292(15): 6339–6351.

[37] Orywal K, Jelski W, Zdrodowski M, et al. The activity of class I, II, III and IV alcohol dehydrogenase isoenzymes and aldehyde dehydrogenase in ovarian cancer and ovarian cysts[J]. Adv Med Sci, 2013, 58(2): 216–20.

[38] Sawers L, Ferguson M J, Ihrig B R, et al. Glutathione S-transferase P1 (GSTP1) directly influences platinum drug chemosensitivity in ovarian tumour cell lines[J]. Br J Cancer, 2014, 111(6): 1150–8.

[39] Keita M, Ainmelk Y, Pelmus M, et al. Endometrioid ovarian cancer and endometriotic cells exhibit the same alteration in the expression of interleukin-1 receptor II: to a link between endometriosis and endometrioid ovarian cancer[J]. J Obstet Gynaecol Res, 2011, 37(2): 99–107.

[40] Chai A W Y, Cheung A K L, Dai W, et al. Elevated levels of serum nidogen-2 in esophageal squamous cell carcinoma[J]. Cancer Biomark, 2018, 21(3): 583–590.

[41] Cotter T P, Kealy N P, Duggan P F, et al. Elevated serum angiotensin converting enzyme levels in metastatic ovarian dysgerminoma[J]. Respir Med, 1997, 91(4): 237–9.

[42] Beyazit F, Ayhan S, Celik H T, et al. Assessment of serum angiotensin-converting enzyme in patients with epithelial ovarian cancer[J]. Arch Gynecol Obstet, 2015, 292(2): 415–20.

[43] Cai G, Ma X, Zou W, et al. Prediction value of intercellular adhesion molecule-1 gene polymorphisms for epithelial ovarian cancer risk, clinical features, and prognosis[J]. Gene, 2014, 546(1): 117–23.

[44] Urunsak I F, Gulec U K, Paydas S, et al. Adenosine deaminase activity in patients with ovarian neoplasms[J]. Arch Gynecol Obstet, 2012, 286(1): 155–9.

[45] Delage B, Fennell D A, Nicholson L, et al. Arginine deprivation and argininosuccinate synthetase expression in the treatment of cancer[J]. Int J Cancer, 2010, 126(12): 2762–72.

[46] Jamieson D, Wilson K, Pridgeon S, et al. NAD(P)H:Quinone Oxidoreductase 1 and NRH:Quinone Oxidoreductase 2 Activity and Expression in Bladder and Ovarian Cancer and Lower NRH:Quinone Oxidoreductase 2 Activity Associated with an NQO2 Exon 3 Single-Nucleotide Polymorphism[J]. Clinical Cancer Research, 2007, 13(5): 1584–1590.

[47] Cortes-Wanstreet M M, Giedzinski E, Limoli C L, et al. Overexpression of glutamate-cysteine ligase protects human COV434 granulosa tumour cells against oxidative and gamma-radiation-induced cell death[J]. Mutagenesis, 2009, 24(3): 211–24.

[48] Hefler-Frischmuth K, Hefler L A, Heinze G, et al. Serum C-reactive protein in the differential diagnosis of ovarian masses[J]. Eur J Obstet Gynecol Reprod Biol, 2009, 147(1): 65–8.

[49] Mandato V D, Magnani E, Abrate M, et al. Haptoglobin phenotype and epithelial ovarian cancer[J]. Anticancer Res, 2012, 32(10): 4353–8.

[50] Haroon S, Idrees R, Fatima S, et al. Ovarian steroid cell tumor, not otherwise specified: a clinicopathological and immunohistochemical experience of 12 cases[J]. J Obstet Gynaecol Res, 2015, 41(3): 424–31.

[51] Yoshida A, Okamoto N, Tozawa-Ono A, et al. Proteomic analysis of differential protein expression by brain metastases of gynecological malignancies[J]. Hum Cell, 2013, 26(2): 56–66.

[52] Gau D M, Lesnock J L, Hood B L, et al. BRCA1 deficiency in ovarian cancer is associated with alteration in expression of several key regulators of cell motility – A proteomics study[J]. Cell Cycle, 2015, 14(12): 1884–92.

[53] Cho M S, Rupaimoole R, Choi H J, et al. Complement Component 3 Is Regulated by TWIST1 and Mediates Epithelial-Mesenchymal Transition[J]. J Immunol, 2016, 196(3): 1412–8.

[54] Gomes J, Gomes-Alves P, Carvalho S B, et al. Extracellular Vesicles from Ovarian Carcinoma Cells Display Specific Glycosignatures[J]. Biomolecules, 2015, 5(3): 1741–61.

[55] Hjerpe E, Egyhazi Brage S, Carlson J, et al. Metabolic markers GAPDH, PKM2, ATP5B and BEC-index in advanced serous ovarian cancer[J]. BMC Clin Pathol, 2013, 13(1): 30.

[56] Deng Y, Chen C, Hua M, et al. Annexin A2 plays a critical role in epithelial ovarian cancer[J]. Arch Gynecol Obstet, 2015, 292(1): 175–82.

[57] Matsuo K, Nishimura M, Komurov K, et al. Platelet-derived growth factor receptor alpha (PDGFRalpha) targeting and relevant biomarkers in ovarian carcinoma[J]. Gynecol Oncol, 2014, 132(1): 166–75.

[58] Torky H A, Sherif A, Abo-Louz A, et al. Evaluation of Serum Nidogen-2 as a Screening and Diagnostic Tool for Ovarian Cancer[J]. Gynecol Obstet Invest, 2018, 83(5): 461–465.

[59] Mehner C, Oberg A L, Kalli K R, et al. Serine protease inhibitor Kazal type 1 (SPINK1) drives proliferation and anoikis resistance in a subset of ovarian cancers[J]. Oncotarget, 2015, 6(34): 35737–54.

[60] Olson S H, Carlson M D, Ostrer H, et al. Genetic variants in SOD2, MPO, and NQO1, and risk of ovarian cancer[J]. Gynecol Oncol, 2004, 93(3): 615–20.

[61] Amano Y, Mandai M, Yamaguchi K, et al. Metabolic alterations caused by HNF1beta expression in ovarian clear cell carcinoma contribute to cell survival[J]. Oncotarget, 2015, 6(28): 26002–17.

[62] Garibay-Cerdenares O L, Hernandez-Ramirez V I, Osorio-Trujillo J C, et al. Haptoglobin and CCR2 receptor expression in ovarian cancer cells that were exposed to ascitic fluid: exploring a new role of haptoglobin in the tumoral microenvironment[J]. Cell Adh Migr, 2015, 9(5): 394–405.

[63] Huang Q, Lan F, Zheng Z, et al. Akt2 kinase suppresses glyceraldehyde-3-phosphate dehydrogenase (GAPDH)-mediated apoptosis in ovarian cancer cells via phosphorylating GAPDH at threonine 237 and decreasing its nuclear translocation[J]. J Biol Chem, 2011, 286(49): 42211–20.

[64] Cai Y Y, Lin W P, Li A P, et al. Combined effects of curcumin and triptolide on an ovarian cancer cell line[J]. Asian Pac J Cancer Prev, 2013, 14(7): 4267–71.

[65] Lim H Y, Ho Q S, Low J, et al. Respiratory competent mitochondria in human ovarian and peritoneal cancer[J]. Mitochondrion, 2011, 11(3): 437–43.

[66] Wang H, Rosen D G, Wang H, et al. Insulin-like growth factor-binding protein 2 and 5 are differentially regulated in ovarian cancer of different histologic types[J]. Mod Pathol, 2006, 19(9): 1149–56.

[67] Cao L, Li X, Zhang Y, et al. Proteomic analysis of human ovarian cancer paclitaxel-resistant cell lines[J]. Zhong Nan Da Xue Xue Bao Yi Xue Ban, 2010, 35(4): 286–94.

[68] Sadacharan S K, Cavanagh A C, Gupta R S. Immunoelectron microscopy provides evidence for the presence of mitochondrial heat shock 10-kDa protein (chaperonin 10) in red blood cells and a variety of secretory granules[J]. Histochem Cell Biol, 2001, 116(6): 507–17.

[69] Luo J, Li Z, Zhu H, et al. A Novel Role of Cab45-G in Mediating Cell Migration in Cancer Cells[J]. Int J Biol Sci, 2016, 12(6): 677–87.

[70] Lim W, Bae S M, Jo G, et al. Prostaglandin D(2) synthase related to estrogen in the female reproductive tract[J]. Biochem Biophys Res Commun, 2015, 456(1): 355–60.

[71] Kryczek I, Lange A, Mottram P, et al. CXCL12 and vascular endothelial growth factor synergistically induce neoangiogenesis in human ovarian cancers[J]. Cancer Res, 2005, 65(2): 465–72.

[72] Nieman K M, Kenny H A, Penicka C V, et al. Adipocytes promote ovarian cancer metastasis and provide energy for rapid tumor growth[J]. Nat Med, 2011, 17(11): 1498–503.

[73] Symowicz J, Adley B P, Gleason K J, et al. Engagement of collagen-binding integrins promotes matrix metalloproteinase-9-dependent E-cadherin ectodomain shedding in ovarian carcinoma cells[J]. Cancer Res, 2007, 67(5): 2030–9.

[74] Levy J M M, Towers C G, Thorburn A. Targeting autophagy in cancer[J]. Nat Rev Cancer, 2017, 17(9): 528–542.

[75] Gao B, Radaeva S, Park O. Liver natural killer and natural killer T cells: immunobiology and emerging roles in liver diseases[J]. J Leukoc Biol, 2009, 86(3): 513–28.

[76] Mak T W, Grusdat M, Duncan G S, et al. Glutathione Primes T Cell Metabolism for Inflammation[J]. Immunity, 2017, 46(4): 675–689.

[77] Waldner M J, Foersch S, Neurath M F. Interleukin-6--a key regulator of colorectal cancer development[J]. Int J Biol Sci, 2012, 8(9): 1248–53.

[78] Andrade K, Fornetti J, Zhao L, et al. RON kinase: A target for treatment of cancer-induced bone destruction and osteoporosis[J]. Sci Transl Med, 2017, 9(374).

[79] Matsuoka T, Yashiro M. Rho/ROCK signaling in motility and metastasis of gastric cancer[J]. World J Gastroenterol, 2014, 20(38): 13756–66.

[80] Thangasamy T, Jeyakumar P, Sittadjody S, et al. L-carnitine mediates protection against DNA damage in lymphocytes of aged rats[J]. Biogerontology, 2009, 10(2): 163–72.

[81] Duran A, Hernandez E D, Reina-Campos M, et al. p62/SQSTM1 by Binding to Vitamin D Receptor Inhibits Hepatic Stellate Cell Activity, Fibrosis, and Liver Cancer[J]. Cancer Cell, 2016, 30(4): 595–609.

[82] Nosov V, Su F, Amneus M, et al. Validation of serum biomarkers for detection of early-stage ovarian cancer[J]. Am J Obstet Gynecol, 2009, 200(6): 639 e1–5.

[83] Hariprasad G, Hariprasad R, Kumar L, et al. Apolipoprotein A1 as a potential biomarker in the ascitic fluid for the differentiation of advanced ovarian cancers[J]. Biomarkers, 2013, 18(6): 532–41.

[84] Siva A, Xin H, Qin F, et al. Immune modulation by melanoma and ovarian tumor cells through expression of the immunosuppressive molecule CD200[J]. Cancer Immunol Immunother, 2008, 57(7): 987–96.

[85] Cochrane D R, Tessier-Cloutier B, Lawrence K M, et al. Clear cell and endometrioid carcinomas: are their differences attributable to distinct cells of origin?[J]. J Pathol, 2017, 243(1): 26–36.

[86] Khandakar B, Mathur S R, Kumar L, et al. Tissue biomarkers in prognostication of serous ovarian cancer following neoadjuvant chemotherapy[J]. Biomed Res Int, 2014, 2014: 401245.

[87] Elg S A, Mayer A R, Carson L F, et al. Alpha-1 acid glycoprotein is an immunosuppressive factor found in ascites from ovaria carcinoma[J]. Cancer, 1997, 80(8): 1448–56.

[88] Abdel-Aleem H, Ahmed A, Sabra A M, et al. Serum alpha L-fucosidase enzyme activity in ovarian and other female genital tract tumors[J]. Int J Gynaecol Obstet, 1996, 55(3): 273–9.

[89] Zhao S, Ma Y, Huang X. Trefoil factor 1 elevates the malignant phenotype of mucinous ovarian cancer cell through Wnt/beta-catenin signaling[J]. Int J Clin Exp Pathol, 2015, 8(9): 10412–9.

[90] Carduner L, Agniel R, Kellouche S, et al. Ovarian cancer ascites-derived vitronectin and fibronectin: Combined purification, molecular features and effects on cell response[J]. Biochimica et Biophysica Acta (BBA) – General Subjects, 2013, 1830(10): 4885–4897.

[91] Leinster D A, Colom B, Whiteford J R, et al. Endothelial cell junctional adhesion molecule C plays a key role in the development of tumors in a murine model of ovarian cancer[J]. FASEB J, 2013, 27(10): 4244–53.

[92] Liu Y, Zheng D, Liu M, et al. Downregulation of glypican-3 expression increases migration, invasion, and tumorigenicity of human ovarian cancer cells[J]. Tumour Biol, 2015, 36(10): 7997–8006.

[93] Saraswat M, Mrudula T, Kumar P U, et al. Overexpression of aldose reductase in human cancer tissues[J]. Med Sci Monit, 2006, 12(12): CR525–529.

[94] Durkes A, Garner M, Juan-Salles C, et al. Immunohistochemical characterization of nonhuman primate ovarian sex cord-stromal tumors[J]. Vet Pathol, 2012, 49(5): 834–8.

[95] Shim J H, Xiao C, Hayden M S, et al. CHMP5 is essential for late endosome function and down-regulation of receptor signaling during mouse embryogenesis[J]. J Cell Biol, 2006, 172(7): 1045–56.

[96] Costanzi E, Urbanelli L, Bellezza I, et al. Hypermethylation contributes to down-regulation of lysosomal beta-hexosaminidase alpha subunit in prostate cancer cells[J]. Biochimie, 2014, 101: 75–82.

[97] Goyne H E, Stone P J, Burnett A F, et al. Ovarian tumor ascites CD14+ cells suppress dendritic cell-activated CD4+ T-cell responses through IL-10 secretion and indoleamine 2,3-dioxygenase[J]. J Immunother, 2014, 37(3): 163–9.

[98] Li K, Dan Z, Hu X, et al. CD14 regulates gastric cancer cell epithelialmesenchymal transition and invasion in vitro[J]. Oncol Rep, 2013, 30(6): 2725–32.

[99] Usui A, Ko S Y, Barengo N, et al. P-cadherin promotes ovarian cancer dissemination through tumor cell aggregation and tumor-peritoneum interactions[J]. Mol Cancer Res, 2014, 12(4): 504–13.

[100] Ayyub A, Saleem M, Fatima I, et al. Glycosylated Alpha-1-acid glycoprotein 1 as a potential lung cancer serum biomarker[J]. Int J Biochem Cell Biol, 2016, 70: 68–75.

[101] Okwan-Duodu D, Landry J, Shen X Z, et al. Angiotensin-converting enzyme and the tumor microenvironment: mechanisms beyond angiogenesis[J]. Am J Physiol Regul Integr Comp Physiol, 2013, 305(3): R205–15.

[102] Tucker S L, Gharpure K, Herbrich S M, et al. Molecular biomarkers of residual disease after surgical debulking of high-grade serous ovarian cancer[J]. Clin Cancer Res, 2014, 20(12): 3280–8.

[103] Odening K E, Li W, Rutz R, et al. Enhanced complement resistance in drug-selected P-glycoprotein expressing multi-drug-resistant ovarian carcinoma cells[J]. Clin Exp Immunol, 2009, 155(2): 239–48.

[104] Lee C S, Taib N A, Ashrafzadeh A, et al. Unmasking Heavily O-Glycosylated Serum Proteins Using Perchloric Acid: Identification of Serum Proteoglycan 4 and Protease C1 Inhibitor as Molecular Indicators for Screening of Breast Cancer[J]. PLoS One, 2016, 11(2): e0149551.

[105] Yamamoto K, Ohga N, Hida Y, et al. Biglycan is a specific marker and an autocrine angiogenic factor of tumour endothelial cells[J]. Br J Cancer, 2012, 106(6): 1214–23.

[106] Yousif N G. Fibronectin promotes migration and invasion of ovarian cancer cells through up-regulation of FAK-PI3K/Akt pathway[J]. Cell Biol Int, 2014, 38(1): 85–91.

[107] Yip P, Chen T H, Seshaiah P, et al. Comprehensive serum profiling for the discovery of epithelial ovarian cancer biomarkers[J]. PLoS One, 2011, 6(12): e29533.

[108] Isgro M A, Bottoni P, Scatena R. Neuron-Specific Enolase as a Biomarker: Biochemical and Clinical Aspects[J]. Adv Exp Med Biol, 2015, 867: 125–43.

[109] Xi H Q, Cai A Z, Wu X S, et al. Leucine-rich repeat-containing G-protein-coupled receptor 5 is associated with invasion, metastasis, and could be a potential therapeutic target in human gastric cancer[J]. Br J Cancer, 2014, 110(8): 2011–20.

[110] Schutte A, Lottaz D, Sterchi E E, et al. Two alpha subunits and one beta subunit of meprin zinc-endopeptidases are differentially expressed in the zebrafish Danio rerio[J]. Biol Chem, 2007, 388(5): 523–31.

[111] Becker-Pauly C, Howel M, Walker T, et al. The alpha and beta subunits of the metalloprotease meprin are expressed in separate layers of human epidermis, revealing different functions in keratinocyte proliferation and differentiation[J]. J Invest Dermatol, 2007, 127(5): 1115–25.

[112] Trivedi S, Starz-Gaiano M. Drosophila Jak/STAT Signaling: Regulation and Relevance in Human Cancer and Metastasis[J]. Int J Mol Sci, 2018, 19(12).

[113] Bonetto A, Aydogdu T, Kunzevitzky N, et al. STAT3 activation in skeletal muscle links muscle wasting and the acute phase response in cancer cachexia[J]. PLoS One, 2011, 6(7): e22538.

[114] Yeung T L, Leung C S, Yip K P, et al. Cellular and molecular processes in ovarian cancer metastasis. A Review in the Theme: Cell and Molecular Processes in Cancer Metastasis[J]. Am J Physiol Cell Physiol, 2015, 309(7): C444–56.

[115] Yang C, Zhou C, Li J, et al. Quantitative proteomic study of the plasma reveals acute phase response and LXR/RXR and FXR/RXR activation in the chronic unpredictable mild stress mouse model of depression[J]. Mol Med Rep, 2018, 17(1): 93–102.

[116] Nowicka A, Marini F C, Solley T N, et al. Human omental-derived adipose stem cells increase ovarian cancer proliferation, migration, and chemoresistance[J]. PLoS One, 2013, 8(12): e81859.

[117] Dirat B, Bochet L, Dabek M, et al. Cancer-associated adipocytes exhibit an activated phenotype and contribute to breast cancer invasion[J]. Cancer Res, 2011, 71(7): 2455–65.

[118] Gee K, Guzzo C, Che Mat N F, et al. The IL-12 family of cytokines in infection, inflammation and autoimmune disorders[J]. Inflamm Allergy Drug Targets, 2009, 8(1): 40–52.

[119] Netea-Maier R T, Smit J W A, Netea M G. Metabolic changes in tumor cells and tumor-associated macrophages: A mutual relationship[J]. Cancer Lett, 2018, 413: 102–109.

[120] Geck R C, Toker A. Nonessential amino acid metabolism in breast cancer[J]. Adv Biol Regul, 2016, 62: 11–17.

[121] Bach J P, Deuster O, Balzer-Geldsetzer M, et al. The role of macrophage inhibitory factor in tumorigenesis and central nervous system tumors[J]. Cancer, 2009, 115(10): 2031–40.

[122] Roggiani F, Mezzanzanica D, Rea K, et al. Guidance of Signaling Activations by Cadherins and Integrins in Epithelial Ovarian Cancer Cells[J]. Int J Mol Sci, 2016, 17(9).

[123] Sarrouilhe D, Clarhaut J, Defamie N, et al. Serotonin and cancer: what is the link?[J]. Curr Mol Med, 2015, 15(1): 62–77.

[124] Nourshargh S, Alon R. Leukocyte migration into inflamed tissues[J]. Immunity, 2014, 41(5): 694–707.

[125] Berraondo P, Minute L, Ajona D, et al. Innate immune mediators in cancer: between defense and resistance[J]. Immunol Rev, 2016, 274(1): 290–306.

[126] Van Der Steen S C, Van Tilborg A A, Vallen M J, et al. Prognostic significance of highly sulfated chondroitin sulfates in ovarian cancer defined by the single chain antibody GD3A11[J]. Gynecol Oncol, 2016, 140(3): 527–36.

[127] Davalieva K, Kiprijanovska S, Maleva Kostovska I, et al. Comparative Proteomics Analysis of Urine Reveals Down-Regulation of Acute Phase Response Signaling and LXR/RXR Activation Pathways in Prostate Cancer[J]. Proteomes, 2017, 6(1).

[128] Suryawanshi S, Huang X, Elishaev E, et al. Complement pathway is frequently altered in endometriosis and endometriosis-associated ovarian cancer[J]. Clin Cancer Res, 2014, 20(23): 6163–74.

[129] Afshar-Kharghan V. The role of the complement system in cancer[J]. J Clin Invest, 2017, 127(3): 780–789.

[130] Bareke H, Akbuga J. Complement system’s role in cancer and its therapeutic potential in ovarian cancer[J]. Scand J Immunol, 2018, 88(1): e12672.

[131] Corrales L, Matson V, Flood B, et al. Innate immune signaling and regulation in cancer immunotherapy[J]. Cell Res, 2017, 27(1): 96–108.

[132] Weber M S, Hemmer B. Cooperation of B cells and T cells in the pathogenesis of multiple sclerosis[J]. Results Probl Cell Differ, 2010, 51: 115–26.

[133] Birrer M J, Johnson M E, Hao K, et al. Whole genome oligonucleotide-based array comparative genomic hybridization analysis identified fibroblast growth factor 1 as a prognostic marker for advanced-stage serous ovarian adenocarcinomas[J]. J Clin Oncol, 2007, 25(16): 2281–7.

[134] Sperner-Unterweger B, Neurauter G, Klieber M, et al. Enhanced tryptophan degradation in patients with ovarian carcinoma correlates with several serum soluble immune activation markers[J]. Immunobiology, 2011, 216(3): 296–301.

[135] Lu J, Tan M, Cai Q. The Warburg effect in tumor progression: mitochondrial oxidative metabolism as an anti-metastasis mechanism[J]. Cancer Lett, 2015, 356(2 Pt A): 156–64.

[136] Wei L, Zhou Y, Yao J, et al. Lactate promotes PGE2 synthesis and gluconeogenesis in monocytes to benefit the growth of inflammation-associated colorectal tumor[J]. Oncotarget, 2015, 6(18): 16198–214.

[137] Shao C, Li M, Li X, et al. A tool for biomarker discovery in the urinary proteome: a manually curated human and animal urine protein biomarker database[J]. Mol Cell Proteomics, 2011, 10(11): M111 010975.

[138] Jing J, Gao Y. Urine biomarkers in the early stages of diseases: current status and perspective[J]. Discov Med, 2018, 25(136): 57–65.

